# Moving toward generalizable NZ-1 labeling for 3D structure determination with optimized epitope tag insertion

**DOI:** 10.1101/2020.10.06.325969

**Authors:** Risako Tamura-Sakaguchi, Rie Aruga, Mika Hirose, Toru Ekimoto, Takuya Miyake, Yohei Hizukuri, Rika Oi, Mika K. Kaneko, Yukinari Kato, Yoshinori Akiyama, Mitsunori Ikeguchi, Kenji Iwasaki, Terukazu Nogi

## Abstract

Antibody labeling has been extensively conducted for structure determination in both x-ray crystallography and EM analysis. However, establishing target-specific antibodies is a prerequisite for applying antibody-assisted structural analysis. To expand the applicability of this strategy, we have developed an alternative method to prepare an antibody-complex by inserting an exogenous epitope into the target. We have already demonstrated that the Fab of monoclonal antibody NZ-1 could form a stable complex with the target containing a PA12 tag as an inserted epitope. Nevertheless, we also found that the complex formation through the inserted PA12 tag inevitably caused structural change around the insertion site of the target. Hence, we here attempted to improve the insertion method and consequently discovered that utilization of a PA14 tag significantly reduced the structural change in the target. By adopting a closed ring-like conformation inside the antigen-binding pocket, the inserted PA14 tag had less impact on the folding of the target. Due to this structural property, the PA14 tag could also be inserted into the sterically hindered loop for labeling. Molecular dynamics simulations also indicated that the folding of the target was rigid regardless of the PA14 insertion and the complex formation with the NZ-1 Fab. Using the improved labeling technique, we performed negative-stain EM on a bacterial site-2 protease, which enabled us to approximate the domain arrangement based on the docking mode of the NZ-1 Fab.

## Introduction

Antibody labeling has become useful tools for structure determination of protein molecules and complexes. It is established that the antigen-binding fragment (Fab) and the variable fragment (Fv) can bind to their target and serve as chaperones to promote crystallization in x-ray crystallography (Hino *et al*., 2013, Hunte & Michel, 2002, Koide, 2009). Accordingly, a large number of crystal structures have been successfully determined using crystallization chaperones for membrane proteins that were difficult to crystallize without chaperones. The first example of antibody labeling was the structure determination of bacterial cytochrome *c* oxidase in complex with an Fv fragment (Ostermeier *et al*., 1995). Subsequently, the high-resolution crystal structure of KcsA K^+^ channel was successfully determined using an Fab as a crystallization chaperone (Zhou *et al*., 2001). Utilizing Fabs has also accelerated the structure determination of G-protein coupled receptors such as the human β_2_ adrenergic receptor (Rasmussen *et al*., 2007, Day *et al*., 2007). As a consequence, the usefulness of antibody labeling has been broadly accepted in structural biology. Nevertheless, establishing antibodies that stably bind the respective targets is a prerequisite for utilizing antibody labeling and limits the applicability.

To make antibody-assisted structural analysis more immediately applicable, we developed an alternative strategy, in which antibody labeling is mediated through an exogenous epitope sequence that is inserted into the target. Specifically, we utilized the PA tag-NZ-1 antibody pair. The high-affinity NZ-1 monoclonal antibody was established by immunizing rats with a tetradecapeptide (EGGVAMPGAEDDVV) from the platelet-aggregation-stimulating domain of human podoplanin (Kato *et al*., 2006). It was also shown that the truncated dodecapeptide (GVAMPGAEDDVV) can bind to NZ-1 with an affinity comparable to that for the original epitope (Fujii *et al*., 2014), and has been developed as PA tag for affinity purification and specific labeling (in this study, PA tag is referred to as PA12 tag while the original epitope composed of 14 residues is referred to as PA14 tag for clarity). The recognition mode was examined by determining the crystal structure of the NZ-1 Fab in complex with the PA14 peptide (Fujii *et al*., 2016). The structure shows that the PA12 portion of PA14 peptide binds stably with NZ-1 and adopts a bent loop-like conformation in the antigen-binding pocket of NZ-1. These observations prompted the prediction that the PA12 tag could bind with NZ-1 either when fused to the termini of the target or when inserted into a loop region. In fact, our previous study demonstrated the preparation of multiple crystallizable complexes with the PA12 tag inserted into loop regions protruding from a globular domain of the target (Tamura *et al*., 2019). Our target protein was an integral membrane protein from a hyperthermophile *Aquifex aeolicus* (Deckert *et al*., 1998). This membrane protein is an orthologue of the *E. coli* intramembrane protease RseP that belongs to the site-2 protease family (Hizukuri *et al*., 2017) (hereafter, *E. coli* RseP and the *A. aeolicus* orthologue are referred to as *Ec*RseP and *Aa*RseP, respectively). Like *Ec*RseP, *Aa*RseP possesses two tandemly-arranged PSD95/Dlg/ZO-1 (PDZ) domains in the periplasmic soluble region, referred to as the PDZ tandem (Fig. 1A). The two PDZ domains in the tandem are composed of six β-strands and two α-helices, respectively (Hizukuri *et al*., 2014). Both of the PDZ domains are circular permutants of the canonical PDZ-fold (Fig. 1B). In the PDZ domains of *Aa*RseP, the new termini are formed by a chain-break between strands βB and βC of the canonical fold, and the canonical termini on the opposite side of the PDZ domain at strands βA and βF are connected by a hairpin loop. We demonstrated that the PA12 tag could be inserted into the βF-βA loops of the two PDZ domains without disrupting their structures and that the PA12-inserted PDZ tandem fragment formed a complex with the NZ-1 Fab (Tamura *et al*., 2019).

**Figure 1.**
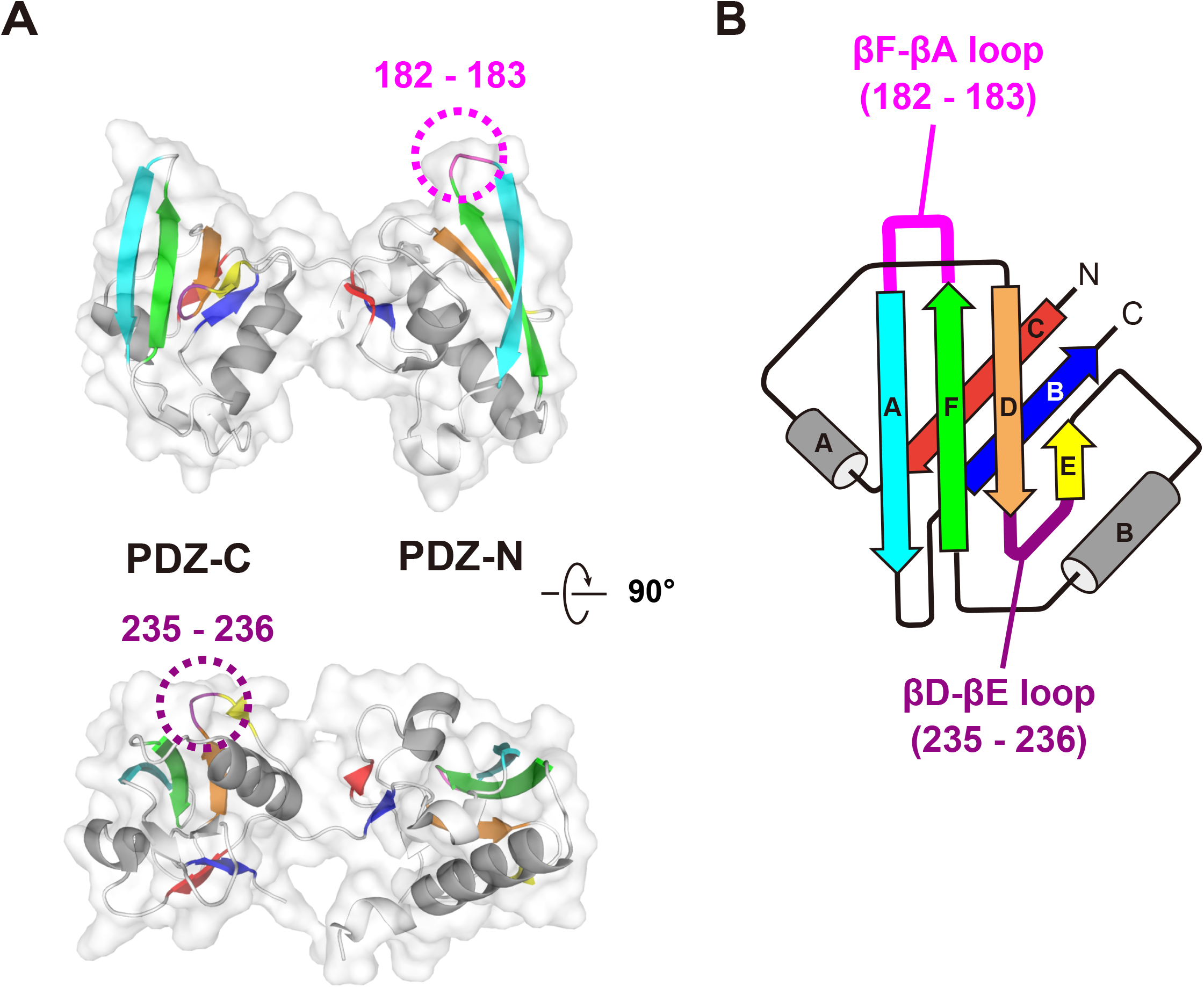
Design of PA14 insertion sites in the PDZ tandem of *Aa*RseP. (A) 3D structure of PDZ tandem of *Aa*RseP. The PDZ tandem structure (PDB ID: 3WKM, chain A) is shown as a ribbon model with transparent surface in two different views. Each PDZ domain is colored by the six β-strands. The α-helices conserved in the PDZ fold are colored gray. The two PA14 insertion sites are highlighted with dotted circles. (B) Topology diagram. Both of the PDZ domains in *Aa*RseP are circular-permutants of the canonical PDZ fold. The first mutant was constructed by replacing residues 182-183 of the βF-βA loop of the PDZ-N domain to the PA14 tag. In the second mutant, PA14 was inserted between residues 235-236 of the βD-βE loop of the PDZ-C domain.

Our previous study also revealed that more rigid complexes could be prepared by adjusting the insertion point to reduce the number of residues that undergo structural change upon binding to the NZ-1 Fab (Tamura *et al*., 2019). However, such residues could not be eliminated completely. For instance, the inter-strand hydrogen bonds of the βF-βA loops were broken around the junctions because the Cα atoms of both ends of PA12 tag were separated by 12-14 Å when accommodated in the antigen-binding pocket of NZ-1. This separation seemed to be inevitable as long as we utilized the PA12 tag as the inserted epitope. Therefore, we here tested if the structural change of the target could be suppressed by modifying the tag sequence in this study. Specifically, we utilized the PA14 tag, instead of PA12, which contains two additional residues (Glu-Gly) upstream of PA12. These two residues were flexible in the co-crystal structure of the NZ-1 Fab with the PA14 peptide, and they may be able to change conformation to accommodate the fold of the target protein near the insertion site. Here we again utilized the PDZ tandem of *Aa*RseP as the target for the PA14 insertion and complex formation with the NZ-1 Fab. In addition to structure determination by x-ray crystallography, we examined the dynamic properties of the PA14-mediated Fab-PDZ complexes through the all-atom molecular dynamics simulations. Finally, we approximated the spatial arrangement of the two PDZ domains in PA14-inserted full-length *Aa*RseP by using negative-stain EM together with our improved NZ-1 labeling technique.

## Results

### Reduction of the structural change of the target by inserting the PA14 tag

We first inserted the PA14 tag between βF and βA in the PDZ-N domain, where we previously inserted the PA12 tag and optimized the linkers (Tamura *et al*., 2019) (Fig. 1). As βF-βA protrudes from the globular PDZ-N domain, it is highly possible that the NZ-1 can bind the inserted PA14 tag without steric hindrance. At the same time, this must be balanced with the need for a stiff linker to reduce conformational flexibility in the complex with the NZ-1 Fab. Hence, we deleted two protruding turn residues (Asn-182 and Gly-183) and inserted the PA14 tag between Arg-181 and Glu-184 (hereafter, this mutant is referred to as PDZ tandem (181-PA14-184)). Similar to the PA12-inserted mutants of PDZ tandem in our previous study, the purified PDZ tandem (181-PA14-184) was stable and monodisperse, indicating that the PA14 insertion did not disrupt the folding of PDZ tandem. The complex of PDZ tandem (181-PA14-184) with the NZ-1 Fab was also stable and produced crystals under several conditions. X-ray diffraction data were collected up to 2.5 Å resolution (Table 1), and we assigned a PDZ tandem and an NZ-1 Fab in the asymmetric unit by molecular replacement. The crystal packing was maintained by both of the two molecules (Fig. 2A, B). The electron densities were clear enough to build reliable models for the both the PDZ tandem (181-PA14-184) and the NZ-1 Fab (Fig. 2C). However, the refined model of the PDZ-C domain showed higher temperature factors probably because this domain made no direct contacts with the complex-forming NZ-1 Fab in the crystal (Table 2).

**Table 1.** Data Collection Statistics of Fab-complexes

| PDZ tandem | 181-PA14-184 | 235-PA14-236 |
| --- | --- | --- |
| Space group | $P2_12_12_1$ | $P2_1$ |
| Cell dimensions |  |  |
| $a, b, c$ (Å) | 52.36, 75.20, 172.53 | 81.36, 80.18, 168.54 |
| $\alpha, \beta, \gamma$ (°) | 90, 90, 90 | 90, 95.7, 90 |
| No. of complexes / a.s.u. | 1 | 2 |
| X-ray source | PF/BL-5A | PF/BL-17A |
| Wavelength (Å) | 1.0000 | 0.9800 |
| Resolution limits (Å) | 45.68-2.50 (2.60-2.50) | 38.99-3.20 (3.36-3.20) |
| No. of unique reflection | 24,452 (2,685) | 35,827 (4,745) |
| Completeness (%) | 99.9 (99.6) | 99.5 (99.7) |
| Redundancy | 6.6 (6.8) | 3.4 (3.5) |
| $I/\sigma(I)$ | 13.1 (1.5) | 8.0 (1.2) |
| $R_{\text{merge}}$ | 0.093 (1.469) | 0.115 (0.965) |
| $R_{\text{meas}}$ | 0.102 (1.589) | 0.137 (1.141) |
| CC (1/2) | 0.998 (0.674) | 0.994 (0.774) |

Values in parentheses are for highest-resolution shell.

**Figure 2.**
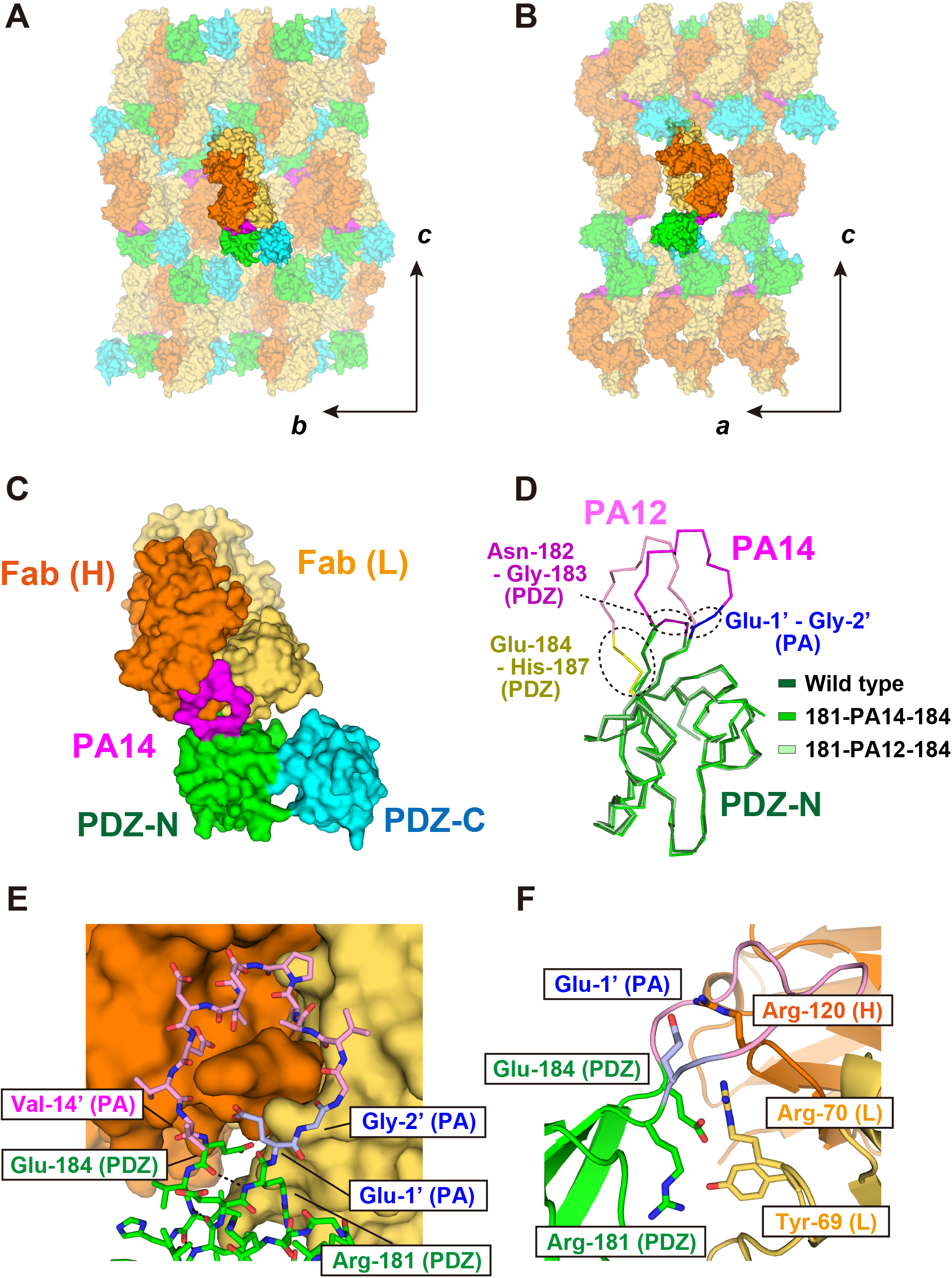
Crystal structure of the PDZ tandem (181-PA14-184) complexed with the NZ-1 Fab. (A, B) Crystal packing in two different views. The crystallographic axes are indicated with arrows. The complex is shown as a surface model and the neighboring complexes in the crystal lattice are shown in partial transparency. The heavy chain and light chain of Fab were colored dark and light orange, respectively. The PDZ-N and -C domains are colored green and cyan, respectively where the inserted PA14 tag is colored magenta. (C) Overall structure of the complex. The NZ-1 Fab docks to PDZ-N through PA14 and does not have direct contacts with PDZ-C within the complex. (D) Superposition of the Cα traces. The wild type PDZ-N domain and the PA12- and PA14-inserted mutants are colored as in (C) and shown in light, medium, and dark colors, respectively. The structures of the wild type and PA12-inserted PDZ-N domains were extracted from the crystal structures determined previously (PDB ID: 3WKL (wild type), 6AL1 (PA12-inserted mutant)). The turn residues replaced by the insertion (Asn-182 and Gly-183) and the inserted sequences (PA12 and PA14) are colored magenta except that Glu-1’ and Gly-2’ on PA 14 are colored blue. In the PA12-inserted mutant, junction residues highlighted in yellow (Glu-184 to His-187) alter their conformations by PA12 insertion and complex formation. In contrast, the addition of Glu-1’ and Gly-2’ seemed to suppress the structural change of the target PDZ-N domain including the junction residues in the PA14-inserted mutant. (E) Close-up view of the binding site. The NZ-1 Fab is shown as in (C). The PDZ-N domain and the inserted PA14 are shown as stick models and colored as in (D). Arg-181 and Glu-184 on the PDZ-N domain maintained the inter-strand hydrogen bond as shown with dotted lines. (F) Residues at the binding interface. The binding interface from (E) is shown as a ribbon model and in profile. The residue whose side chains presumed to form salt bridges or hydrogen bonds are displayed as stick models.

**Table 2.**
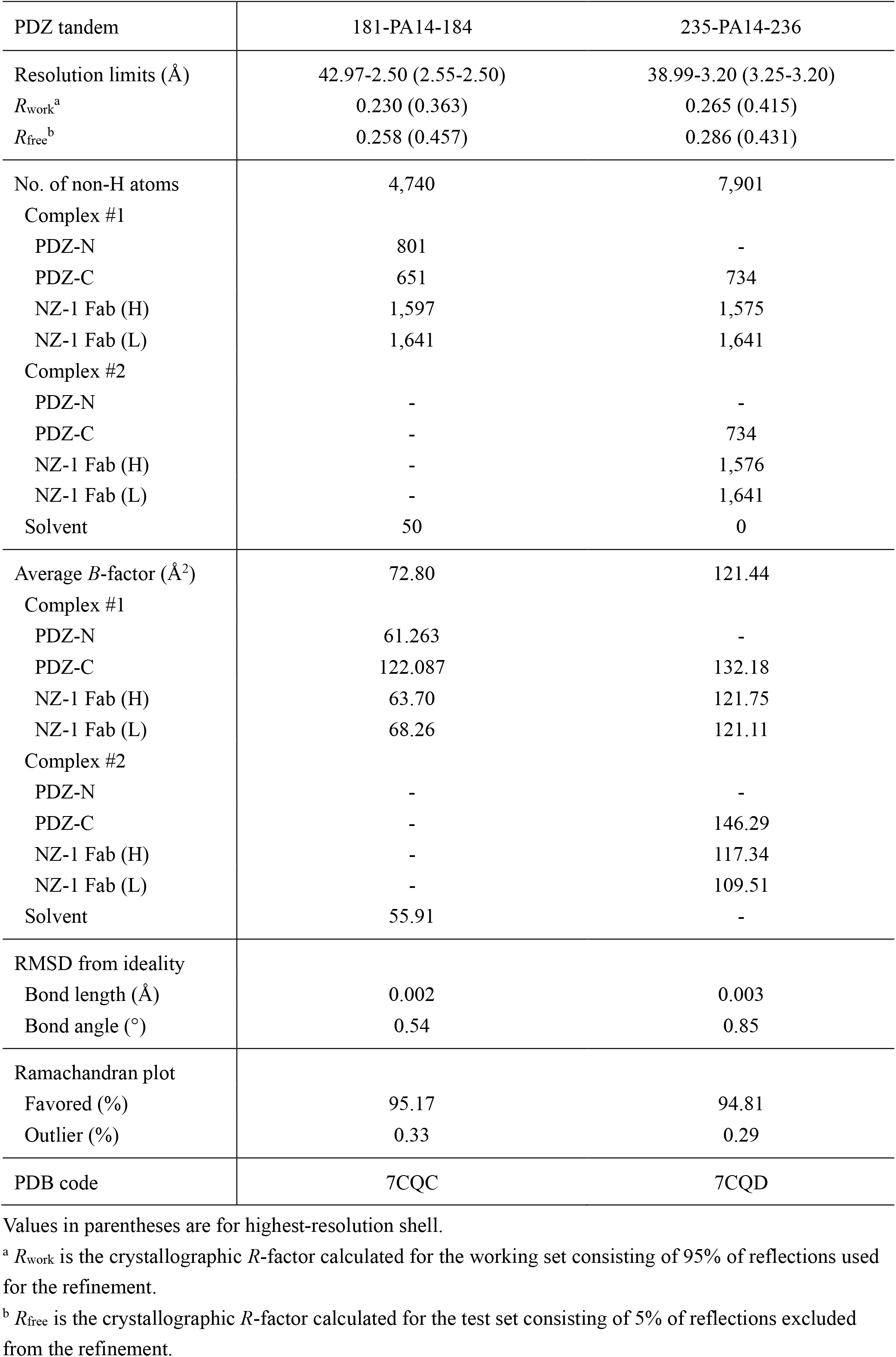
Refinement Statistics of Fab-complexes

| PDZ tandem | 181-PA14-184 | 235-PA14-236 |
| --- | --- | --- |
| Resolution limits (Å) | 42.97-2.50 (2.55-2.50) | 38.99-3.20 (3.25-3.20) |
| $R_{\text{work}}^{\text{a}}$ | 0.230 (0.363) | 0.265 (0.415) |
| $R_{\text{free}}^{\text{b}}$ | 0.258 (0.457) | 0.286 (0.431) |
| No. of non-H atoms | 4,740 | 7,901 |
| Complex #1 |  |  |
| PDZ-N | 801 | - |
| PDZ-C | 651 | 734 |
| NZ-1 Fab (H) | 1,597 | 1,575 |
| NZ-1 Fab (L) | 1,641 | 1,641 |
| Complex #2 |  |  |
| PDZ-N | - | - |
| PDZ-C | - | 734 |
| NZ-1 Fab (H) | - | 1,576 |
| NZ-1 Fab (L) | - | 1,641 |
| Solvent | 50 | 0 |
| Average $B$-factor (Å<sup>2</sup>) | 72.80 | 121.44 |
| Complex #1 |  |  |
| PDZ-N | 61.263 | - |
| PDZ-C | 122.087 | 132.18 |
| NZ-1 Fab (H) | 63.70 | 121.75 |
| NZ-1 Fab (L) | 68.26 | 121.11 |
| Complex #2 |  |  |
| PDZ-N | - | - |
| PDZ-C | - | 146.29 |
| NZ-1 Fab (H) | - | 117.34 |
| NZ-1 Fab (L) | - | 109.51 |
| Solvent | 55.91 | - |
| RMSD from ideality |  |  |
| Bond length (Å) | 0.002 | 0.003 |
| Bond angle (°) | 0.54 | 0.85 |
| Ramachandran plot |  |  |
| Favored (%) | 95.17 | 94.81 |
| Outlier (%) | 0.33 | 0.29 |
| PDB code | 7CQC | 7CQD |

Values in parentheses are for highest-resolution shell.
<sup>a</sup> $R_{\text{work}}$ is the crystallographic $R$ -factor calculated for the working set consisting of 95% of reflections used for the refinement.
<sup>b</sup> $R_{\text{free}}$ is the crystallographic $R$ -factor calculated for the test set consisting of 5% of reflections excluded from the refinement.

As we anticipated, the folding of PDZ-N was not disrupted by the PA14 insertion and complex formation with the NZ-1 Fab where the inserted PA14 tag assumed a closed ring-like structure inside the antigen-binding pocket (Fig. 2D, E). The 80 Cα atoms of PDZ-N, including Arg-181 and Glu-184, were superposed onto those of the wild type with a root-mean-square deviation (RMSD) of 0.750 Å. Arg-181 and Glu-184 at the junctions formed inter-strand hydrogen bonds as is formed in the wild-type PDZ-N (Fig. 2E). The distance between the Cα atoms of Glu-1’ and Val-14’ (hereafter, the residue number in the PA14 tag is indicated with a prime mark) at both ends of the PA14 tag was less than 7 Å, which was remarkably shorter than the separation caused by the PA12 insertion (Fig. 2D). These observations suggest that the NZ-1 labeling with PA14 generally has less impact on the folding of the target than that with PA12.

Unexpectedly, the newly introduced Glu-1’ residue formed a salt-bridge with Arg-120 on the NZ-1 heavy chain (Fig. 2F), whereas the same residue was highly mobile in the co-crystal structure of the NZ-1 Fab with the PA14 peptide (Fujii *et al*., 2016). Besides the inserted PA14 residues, only the junction residues in the PDZ tandem (181-PA14-184) are involved in the inter-molecular interactions with the NZ-1 Fab (Fig. 2F). Arg-181 on PDZ-N at the N-terminal side formed a hydrogen bond with Tyr-69 on the light chain, and Glu-184 on PDZ-N is positioned to form a salt bridge with Arg-70 on the light chain.

### Insertion of the PA14 tag into a sterically hindered loop region

We next asked if the PA14 tag can be successfully inserted into multiple sites such that the PA14 insertion can serve as a generalizable strategy for structure determination. The insertion site for PDZ tandem (181-PA14-184) was chosen because that loop projects into solvent, but the context and orientation of the insertion site will be unknown in many practical applications of this method. To determine if the PA14 tag could be inserted into a more sterically hindered site, we selected the βD-βE loop as the insertion site (Fig. 1). The βD and βE strands belong to a four-stranded β-sheet, and the turn residues (Asn-235 and Gly-236) also contribute to the flat surface of the β-sheet. In contrast to NZ-1 labeling on a protruding loop, steric hindrance between NZ-1 and the target PDZ domain needs to be avoided in this case. Therefore, we inserted the PA14 tag between the turn residues without deletion, resulting in the mutant PDZ tandem (235-PA14-236). The complex of PDZ tandem (235-PA14-236) with the NZ-1 Fab was also successfully crystallized and diffraction data were collected up to 3.2 Å resolution (Table 1). We assigned two complexes (referred to as complex #1 and #2) in the asymmetric unit by molecular replacement, but the electron densities for the PDZ-N domains were too weak to assign reliable models for this domain in either of the two complexes. Judging from the weak electron densities, the PDZ-N domains in both of the two complexes were exposed to the solvent and made little contribution to the lattice formation in the crystal. Hence, the structures of the NZ-1 Fabs and the PA14-inserted PDZ-C domains were refined and included in the final model (Fig 3A, B, and Table 2).

**Figure 3.**
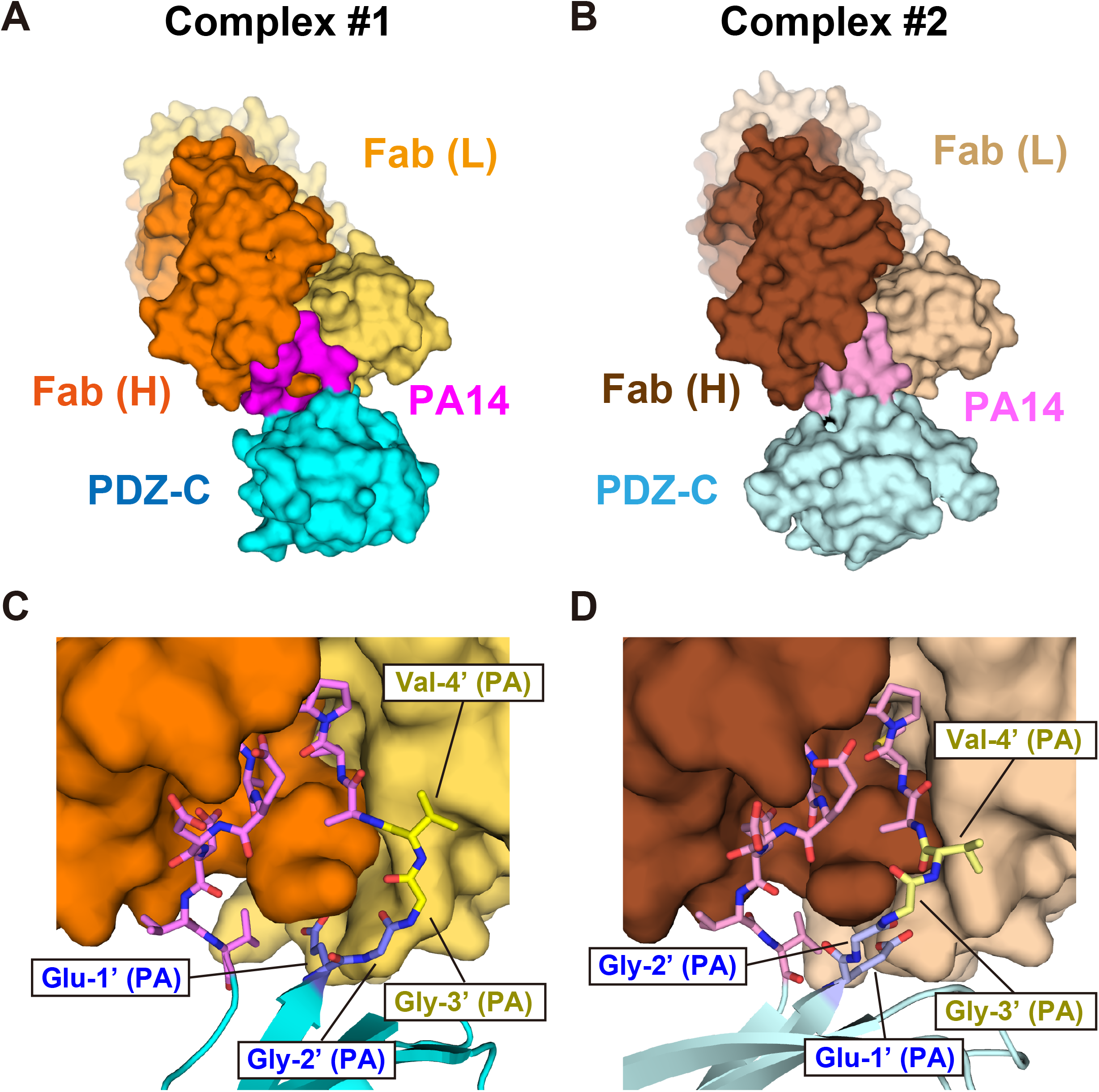
Crystal structure of the PDZ tandem (235-PA14-236) complexed with the NZ-1 Fab. (A, B) The two complexes in the asymmetric unit, complex #1 and #2, are shown as surface models. The heavy and light chains of NZ-1 Fab in complex #1 are colored dark and light orange, respectively while those in complex #2 are colored dark and light brown, respectively. The inserted PA14 tags are colored dark and light magenta, respectively. Due to the disorder in the electron density, the PDZ-N domains were not included in the final model. (C, D) Close-up view of the binding site. The PDZ-C domain and the inserted PA14 residues are shown as ribbon and stick models, respectively. Glu-1’-Gly-2’ and Gly-3’-Val-4’ of PA14 are colored blue and yellow, respectively. The remaining C-terminal ten residues (Ala-5’ to Val-14’) are colored magenta. The residues in complex #1 and #2 are shown in dark and light colors, respectively.

As in PDZ tandem (181-PA14-184), the PA14 tag adopts a closed ring-like structures in both of the two complexes in the asymmetric unit. However, the conformations of the N-terminal residues were significantly different between them (Fig. 3C, D). In all of the crystal structures that have been determined so far, Gly-3’ and Val-4’ on PA14 were located in proximity to Tyr-52 and Tyr-113 on the NZ-1 light chain. In complex #1, the conformations of Gly-3’ and Val-4’ were consistent with those in the known structures, whereas the two residues were largely separated from the NZ-1 light chain in complex #2. It appeared that the conformational change of these two residues was related to crystal packing of the target PDZ-C domains (Fig. 4A, B). In complex #2, the PDZ-C domain intimately contacted the neighboring Fab, resulting in a rigid body reorientation of the PDZ-C domain relative to the complex-forming NZ-1 Fab. Despite the different interaction modes with the NZ-1 Fab, the main chain structures of the PDZ-C domains in both complexes were consistent with that in the wild-type structure without the PA14 insertion. The 81 Cα atoms of PDZ-C, including junction residues, Gly-235 and Asn-236, were superposed onto to those of the wild type with an RMSDs of 0.785 Å and 1.055 Å for complexes #1 and #2, respectively (Fig. 4C). In contrast, the superposition of the Fv region between the two complexes showed the rotation of PDZ-C domain by approximately 40° around the PA14 insertion site (Fig. 4D), which was caused by the conformational change of Gly-3’ and Val-4’. It was previously reported that the replacement of either Gly-3’ or Val-4’ to alanine had no significant effect on the affinity of PA12 tag to NZ-1 (Fujii *et al*., 2014). Presumably, these two residues tolerate conformational change to some extent as their side chains do not have a measurable energetic contribution to the binding affinity. Overall, these observations indicated that the PA14 tag could be inserted into this sterically hindered loop with less structural changes in the target PDZ-C domain.

**Figure 4.**
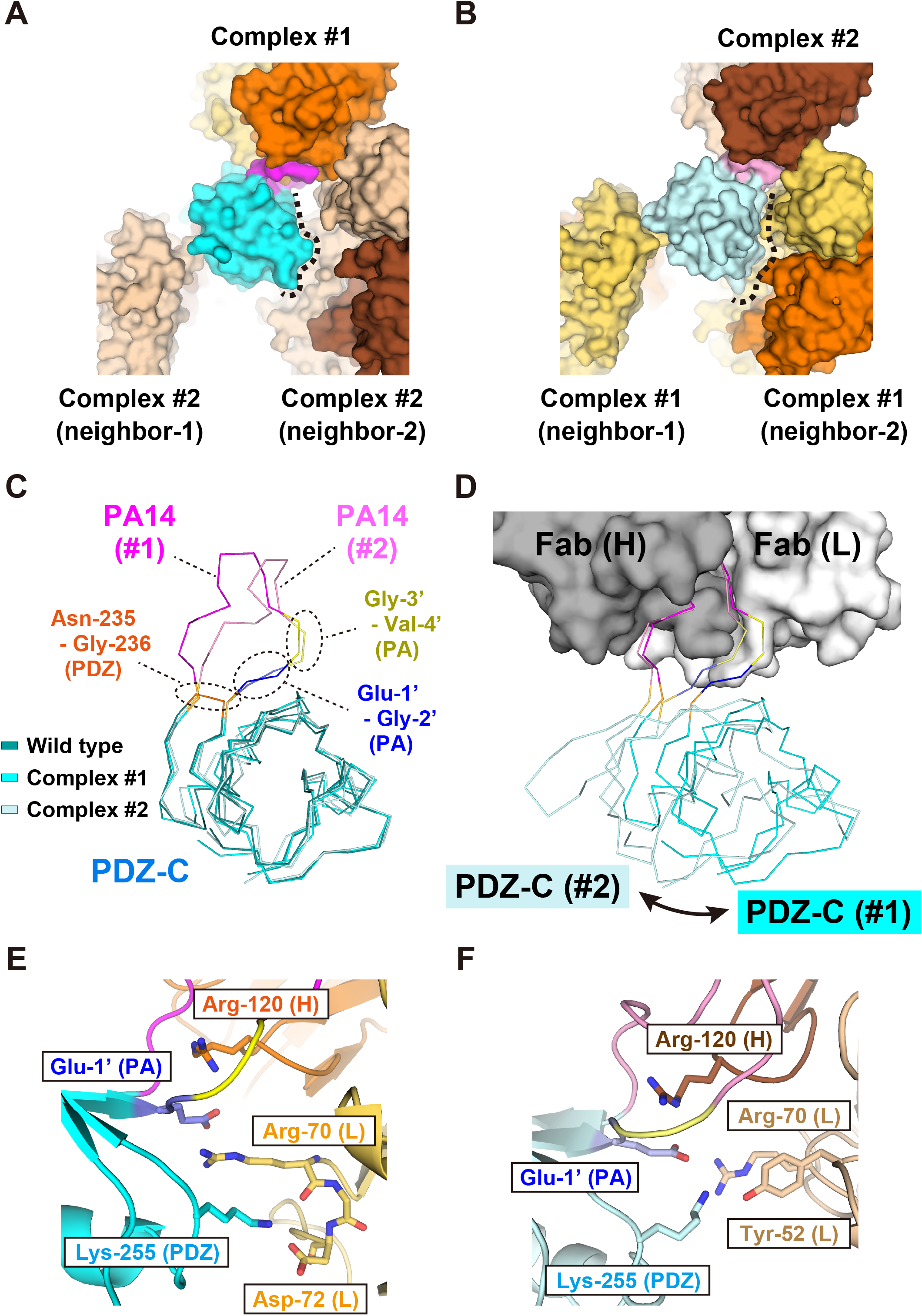
Comparison of the two conformations of the PDZ tandem (235-PA14-236) complexed with the NZ-1 Fab. (A, B) Comparison of crystal contacts in complexes #1 and #2 with PA14-inserted PDZ-C domains. The structures are shown as in Fig. 3A, B. In both complexes #1 and #2, the PDZ-C domains were in direct contact with two neighboring complexes, labeled as neighbor-1 and −2. The PDZ-C domain of complex #1 had a relatively small contact area with neighbor-2 where solvent-accessible space was present between them. Apparently, the relative arrangement of the NZ-1 Fab and PDZ-C in complex #1 was not affected by the crystal packing. In contrast, the PDZ-C domain of complex #2 intimately interacted with neighbor-2, indicating that the positioning of the PDZ-C domain with respect to the NZ-1 Fab was restricted by the crystal packing. (C) Superposition of the Cα traces. The wild type PDZ-C domain (PDB ID: 3WKL) and the PA14-inserted mutants in complexes #1 and #2 are shown in dark, medium, and light colors, respectively. The PDZ-C domain is shown in cyan while the insertion site (Asn-235 and Gly-236) is colored orange. The inserted PA14 tags are colored as in Fig. 3C, D. (D) Superposition of the two complexes based on the NZ-1 Fab. The NZ-1 Fab in complex #1 is shown as a surface model. The heavy chain and light chains are colored grey and white, respectively. The interaction modes of Gly-3’ and Val-4’ of PA14 shown in yellow were different between the two complexes. As a result, the superposition showed a rigid-body rotational movement of the PDZ-C domain around the PA14-insertion site. (E, F) Residues at the binding interfaces of complexes #1 (E) and #2 (F). Residues presumed to form salt bridges or hydrogen bonds are displayed as stick models in the ribbon models.

Concerning the specific interactions between the NZ-1 Fab and PDZ-C, Glu-1’ formed salt bridges with Arg-70 on the NZ-1 light chain as well as to Arg-120 on the heavy chain in both complex #1 and #2 (Fig. 4E, F). In addition to the PA14 residues, Lys-255 on PDZ-C also formed inter-molecular contacts with the NZ-1 Fab, although Lys-255 made different contacts in the two complexes. In complex #1, the amino group of Lys-255 formed hydrogen bonds with the carbonyl group of Arg-70 on the light chain as well as the side chain carboxyl group of Asp-72 on the light chain. In complex #2, the same amino group formed a hydrogen bond with the hydroxyl group of Tyr-52 on the light chain. Thus, the interaction mode of Lys-255 is also affected by the reorientation of PDZ-C relative to the NZ-1 Fab.

### Molecular dynamics simulations to analyze conformational variation in the PA14-mediated Fab-PDZ complexes

For the broadest application of this method, a fixed conformation of the Fab is desirable. Because crystallization may artificially fix the conformation of the complex subunits, we turned to molecular dynamics (MD) simulations to characterize the complex stability and intermolecular motions in the NZ-1 Fab-PDZ complexes. For all complex structures, only the PDZ domain that contains the PA14 insertion was included in the complex model. For the 235-PA14-236 mutant, the structures of both complexes #1 and #2 were used for unique MD simulations. During all simulations, the inserted PA14 tag maintained the closed ring-like conformation inside the antigen-binding pocket, and the PDZ domains did not dissociate from the NZ-1 Fab at all (Supplementary Fig. 1). In all cases, the structures of the PA14-inserted PDZ domains were relatively rigid over 1 μs of simulation, and the RMSDs relative to the initial models were ∼ 2 Å excluding the inserted PA14 residues (Fig. 5A). The results of the simulations suggested that the impact of the PA14 insertion on the folding of the target PDZ domain was relatively small. Focusing on the inserted PA14 tag, the four N-terminal residues, Glu-1’ to Val-4’, showed relatively high root mean square fluctuation (RMSF) (Fig. 5B), which coincides with the observation that the conformation of Gly-3’ and Val-4’ was dependent on the crystal packing in the PDZ tandem (235-PA14-236) with NZ-1 Fab (Fig. 3C, D). With respect to subunit arrangement in the complex, PDZ-N (181-PA14-184) complexed with the NZ-1 Fab was more flexible even though there was only one conformation in the asymmetric unit of the crystal structure. PDZ-N (181-PA14-184) showed higher RMSD and RMSF values than did PDZ-C (235-PA14-236) in the simulations where the structures were aligned at the position of the variable region of the heavy chain (V_H_ region) in the complex (Fig. 5C, E, F, and Supplementary Fig. 2). At 358 ns in the 1-μs simulation, PDZ-N (181-PA14-184) showed the highest RMSD value (∼ 19 Å) relative to the initial model (Supplementary Figs. 1A, 2). In fact, almost no hydrogen-bonding pairs were observed between the NZ-1 Fab and the PDZ-N domain other than those formed by the inserted PA14 tag during the simulation. Additionally, Glu-1’ on PA14 became separated from the NZ-1 Fab whereas this residue was placed close to Arg-120 on the heavy chain in the crystal structure (Fig. 5D). For PDZ-C (235-PA14-236) complexed with the NZ-1 Fab, conformations were preferentially more similar to complex #1 than to complex #2 (Fig. 5E, F). In the simulation initialized from complex #2, the model immediately sampled conformations more similar to complex #1 although conformations similar to complex #2 also appeared as a minor population (Fig. 5F). Comparing the snapshot structures, we found that Lys-255 on PDZ-C formed hydrogen bonds with the residues on the NZ-1 light chain such as Asp-72 in the conformation similar to complex #1 (Supplementary Fig. 3A). In contrast, Lys-255 were separated from Asp-72 in the conformations similar to complex #2 (Supplementary Fig. 3B). Furthermore, the contribution of Glu-1’ on PA14 to the complex formation depended on the PA insertion sites. Glu-1’ in PDZ-C (235-PA14-236) formed a hydrogen bond with Arg-70 on the NZ-1 heavy chain in 72.6% of the snapshots on the trajectory whereas Glu-1’ in PDZ-N (181-PA14-184) reoriented into solvent and away from the interaction with Arg-120 on the NZ-1 heavy chain in the MD simulation as mentioned above.

**Figure 5.**
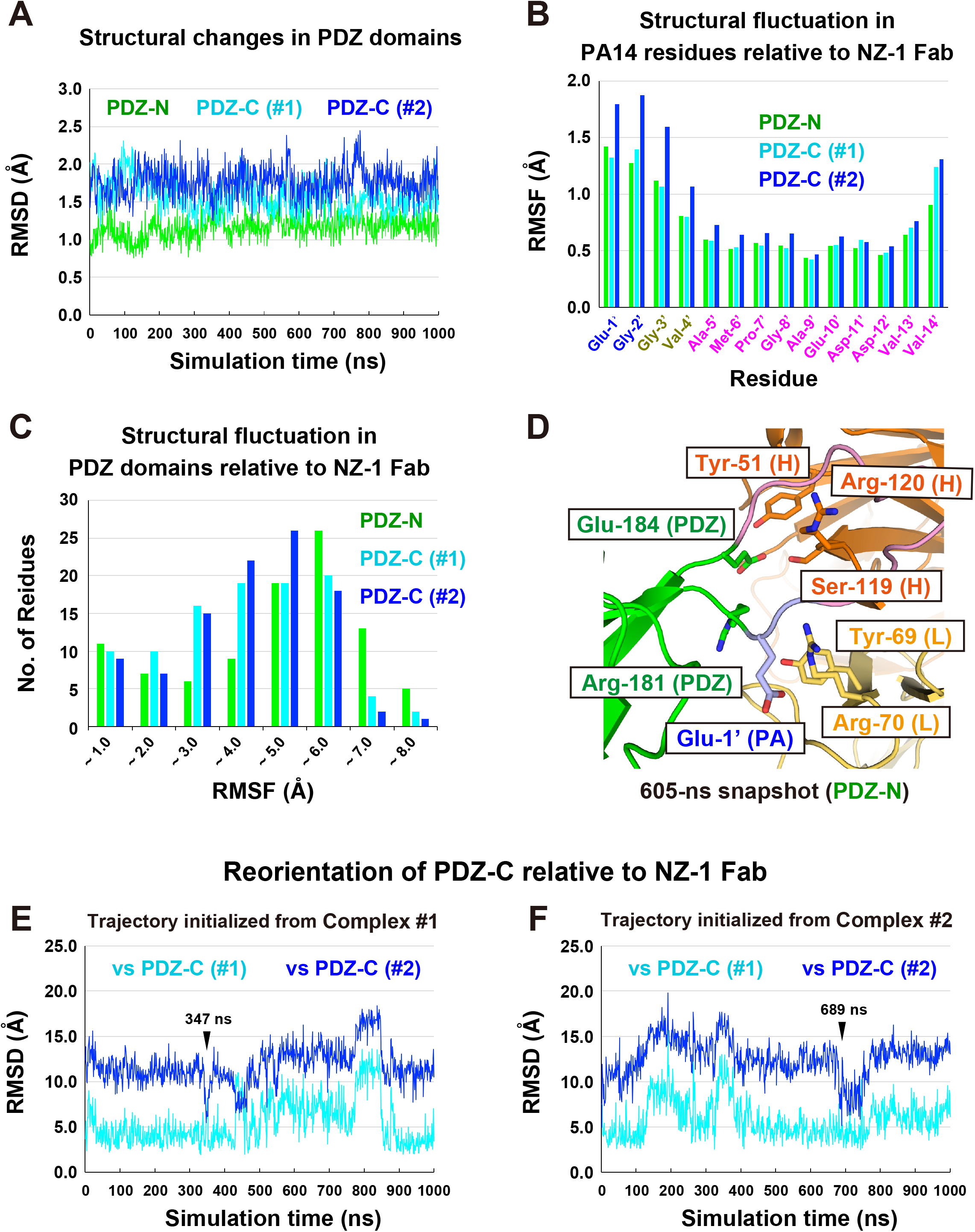
MD simulation of the NZ-1 Fab-PDZ complexes. (A) Trajectories for the target PDZ structures. RMSDs were calculated for the snapshot models relative to the initial model to estimate the structural changes in the PDZ domains during the simulation trajectories. The inserted PA14 residues and the flexible N-terminal region upstream of the PDZ-N domain were excluded from the calculation. RMSDs are plotted for trajectories of the PDZ-N domain (green) and for the PDZ-C domain in both complex #1 (cyan) and #2 (blue). (B) Structural fluctuations in the residues within the PA14 tag. In the RMSF calculations, the complex models within each trajectory were aligned to the initial model based on the V_H_ region of the NZ-1 Fab, and the structural fluctuation from the averaged structure was calculated for each residue on PA14. (C) Histogram of PDZ domain residues binned by RMSF values calculated relative to the NZ-1 Fab. Similar to the process for calculating RMSF values in panel (B), the snapshot models within a trajectory were aligned to the averaged structure at the respective V_H_ regions for each of the three complexes. Subsequently, the fluctuation in the atomic coordinates was calculated for each residue. (D) The representative snapshot of the PDZ-N (181-PA14-184) complexed with NZ-1 Fab from the trajectory. The snapshot at 605 ns, representing the most frequently observed conformation, is shown as a ribbon model. The PDZ-N domain is colored green. PA14 residues are colored light magenta, except for Glu-1’ and Gly-2’ on the N-terminus, which are colored light blue. The heavy and light chains in NZ-1 are colored dark and light orange, respectively. Glu-1’ on PA14 was separated both from Arg-120 on the NZ-1 heavy chain and from Arg-70 on the NZ-1 light chain. Arg-181 was also separated from Tyr-69 on the light chain. Glu-184 on PDZ-N interacted with Tyr-51 and Ser-119 on the NZ-1 heavy chain as opposed to the interaction with Arg-70 on the NZ-1 light chain in the crystal structure (see Fig. 2F). (E, F) Reorientation of the PDZ-C domains relative to the NZ-1 Fab in the MD simulations initialized from complexes #1 (E) and #2 (F), respectively. For both two calculations, the snapshot models were aligned to the initial models of complexes #1 and #2 based on the V_H_ region, and the RMSDs for the PDZ-C domains were calculated relative to the initial models. The RMSDs calculated using complexes #1 and #2 as references are shown in cyan and blue, respectively. The conformations of the 347-ns snapshot in panel (E) and the 689-ns snapshot in panel (F) are similar to that of complex #2, as analyzed in Supplementary Fig. 5C.

### Preparation of the PA14-inserted full-length *Aa*RseP for NZ-1 labeling

Using our improved NZ-1 labeling technique, we next attempted to determine the location and orientation of the two PDZ domains in the context of the full-length *Aa*RseP using antibody-assisted negative-stain EM. Although high-resolution 3D structural data of any full-length RseP orthologue have not yet been elucidated, we previously estimated the spatial arrangement of the PDZ domains using biochemical methods (Hizukuri *et al*., 2014). Specifically, we performed chemical modification analysis to estimate the solvent-accessibility of the surface residues for *Ec*RseP based on the structural data of the PDZ tandem fragment from x-ray crystallography and small-angle scattering analysis. Those previous results suggested that the two PDZ domains adopted a clam-like structure, and that the putative ligand-binding grooves in the PDZ domains would be inaccessible in the context of the full-length protein on the cell membrane. Based on this estimation, the two PA14-insertion sites tested in the present study were presumed to project into solvent because they were located at the back side of the putative ligand-binding grooves. Therefore, it was expected that the PA14 insertion did not destabilized the structure of the full-length protein in both mutants.

We therefore performed negative-stain EM structural analysis on full-length *Aa*RseP constructs with PA14 tag insertions. For these full-length constructs, we used the same insertion sites as used for the PDZ tandem fragments, and produced *Aa*RseP (181-PA14-184) and (235-PA14-236), for PDZ-N and PDZ-C, respectively. After confirming that both mutants accumulated in the *E. coli* membrane, we attempted to examine whether or not the PA14-insertion affected proteolytic activity in *Aa*RseP. However, no physiological substrates have not yet been identified for *Aa*RseP so far. It is known that *Ec*RseP cleaves anti-sigma factor, RseA (Alba *et al*., 2002, Kanehara *et al*., 2002), but no orthologue of RseA has been identified in the *A. aeolicus* genome. Looking for an alternative, we discovered that *Aa*RseP is able to cleave a model substrate for *Ec*RseP derived from RseA (Fig. 6A). In this model substrate, the periplasmic domain of RseA is fused to the first transmembrane region of lactose permease (LY1) with a recombinant cytoplasmic domain containing hemagglutinin (HA)-tagged maltose-binding protein (MBP). Furthermore, the C-terminal periplasmic region of the model substrate was truncated at Val-148 because *Ec*RseP performs the intramembrane proteolysis only after the periplasmic region of RseA is first cleaved by DegS (Alba *et al*., 2002, Kanehara *et al*., 2002). Hence, the model substrate mimics the DegS-cleaved product and can be detected by anti-HA antibody labeling. This model substrate is referred to as HA-MBP-RseA(LY1)148 (Hizukuri & Akiyama, 2012). When co-expressed with either wild-type *Aa*RseP or *Ec*RseP, the band for HA-MBP-RseA(LY1)148 shifts compared with the vector control containing neither *Aa*RseP nor *Ec*RseP. We also observed that the band for HA-MBP-RseA(LY1)148 does not shift when the catalytically important glutamate residue of the “HEXXH” sequence conserved among the site-2 protease family of intramembrane proteases was mutated to glutamine in either *Ec*RseP or *Aa*RseP, which is consistent with intramembrane proteolysis of HA-MBP-RseA(LY1)148 by the respective RsePs. These observations confirm for the first time that *Aa*RseP is functionally orthologous to *Ec*RseP and belongs to the site-2 protease family. Finally, both *Aa*RseP (181-PA14-184) and (235-PA14-236) were also able to cleave HA-MBP-RseA(LY1)148 at comparable levels to the wild-type *Aa*RseP (Fig. 6B), indicating that the PA14-insertion did not abrogate proteolytic activity.

**Figure 6.**
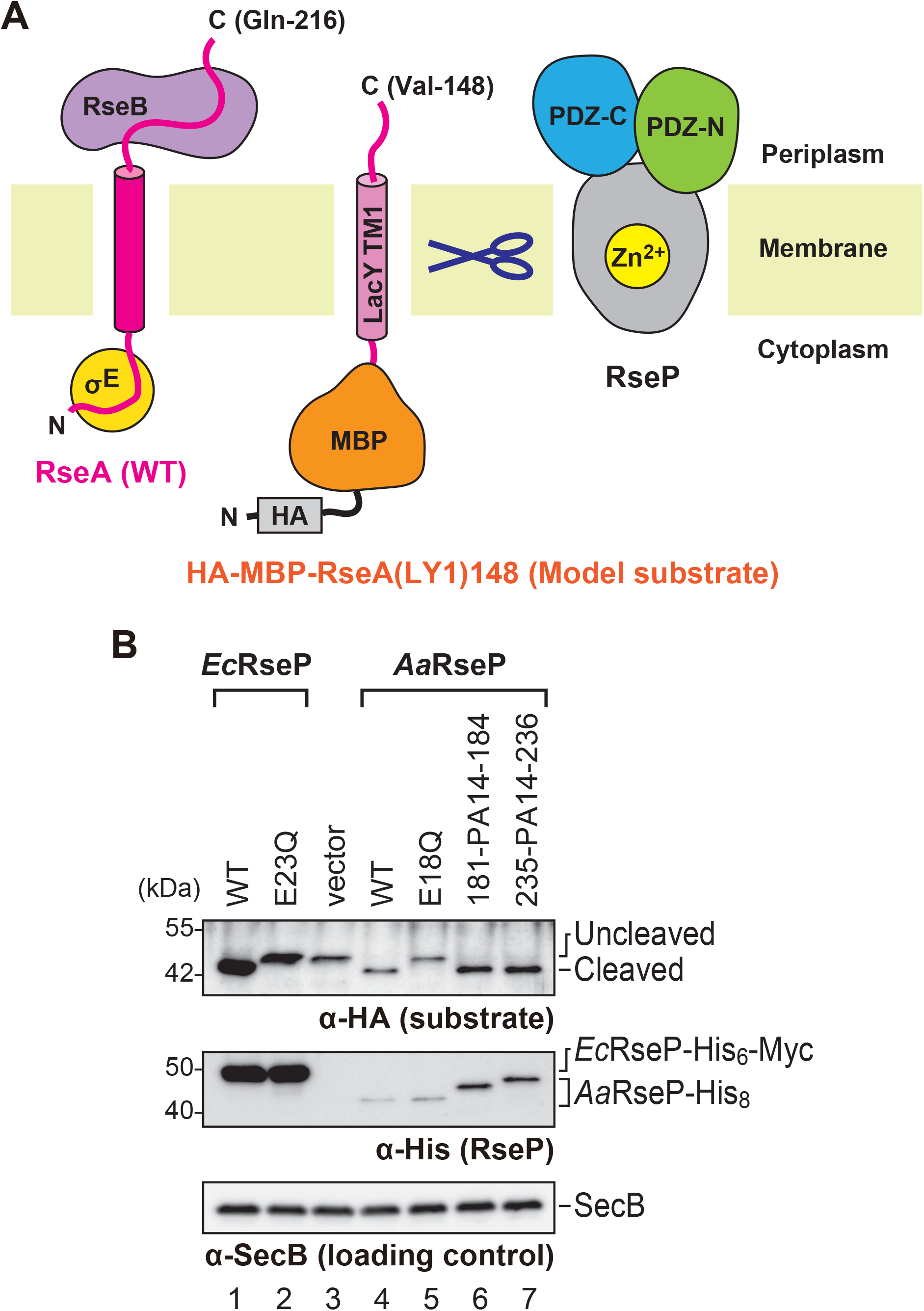
*In vivo* cleavage of model substrate by wild-type *Aa*RseP and PA inserted mutants. (A) Design of the model substrate. The wild type RseA from *E. coli* is a type II membrane protein composed of 216 amino acid residues. The full-length RseA binds with σ^E^ and RseB in the cytoplasmic and periplasmic domains, respectively. DegS cleaves RseA at the C-terminal side of Val-148 and liberates RseB. The model substrate, HA-MBP-RseA(LY1)148, contains the first transmembrane region of lactose permease (LacY TM1: LY1) with a recombinant cytoplasmic domain composed of HA-tagged MBP. The C-terminal periplasmic domain was truncated at Val-148 to mimic the DegS-cleaved product. (B) Immunoblotting with anti-HA antibody detected the model substrate. The full-length model substrate appeared at the *Uncleaved* position. *Cleaved* indicates the model substrate that was cleaved by RseP within the membrane, as illustrated in (A). Immunoblotting with anti-His antibody detected His-tagged *Ec*RseP, *Aa*RseP, and their derivatives. Immunoblotting of cytoplasmic protein SecB serves as a loading control. Molecular size marker positions are shown in kDa on the left.

### Antibody-assisted negative-stain EM of the full-length *Aa*RseP

We next overproduced the PA14-inserted *Aa*RseP mutants in *E. coli* and purified them using immobilized-metal affinity and size-exclusion chromatography. We fractionated monodispersed mutants solubilized with glycol-diosgenin (GDN) detergent, and mixed them with the NZ-1 Fab. We then subjected the mixture to size-exclusion chromatography again to fractionate the stable complex. Subsequently, the purified complex was negatively stained with ammonium molybdate for EM single particle analysis. For *Aa*RseP (181-PA14-184) complexed with the NZ-1 Fab, four different classes of 2D averages were reconstructed into 3D models (Fig. 7A). In each model, the putative Fab fragment was identified by a characteristic ellipsoidal shape with a hole at the center. The remaining part is constituted by a sphere and two protrusions, and presumably corresponds to the detergent-solubilized TM domain and the two PDZ domains of *Aa*RseP. Structural alignment of the four 3D models by the putative *Aa*RseP molecule showed that relative orientation of the NZ-1 Fab among the models hinges around a fixed point (Fig. 7B). This hinge-like variation indicates that the contacting points, namely, the location of the PA14 insertion sites are consistent among the four classes despite the variable orientation of the NZ-1 Fab. It was therefore highly possible that the arrangement of the PDZ-N domain was also consistent among the four classes. In fact, the above-mentioned MD simulation also demonstrated that the orientation of the NZ-1 Fab relative to the PDZ-N (181-PA14-184) fluctuated while the folding of the target PDZ-N domain was maintained throughout the simulation time. For *Aa*RseP (235-PA14-236) complexed with the NZ-1 Fab, the 3D reconstruction also produced four different classes. All of the four 3D models showed similar conformations while the EM maps of three classes were disordered (Supplementary Fig. 4). This observation is also consistent with the result from the MD simulation where the orientation of the NZ-1 Fab relative to the PDZ-C (235-PA14-236) was more fixed as compared to the PDZ-N (181-PA14-184). In the most well-averaged class, the distinctive shape of the NZ-1 Fab was recognized while the two protrusions were again identified on the spherical part (Fig. 7C). Taken together, we could successfully obtain the 3D structural data of the full-length RseP orthologue for the first time, which could be utilized to analyze the spatial arrangement of the two PDZ domains.

**Figure 7.**
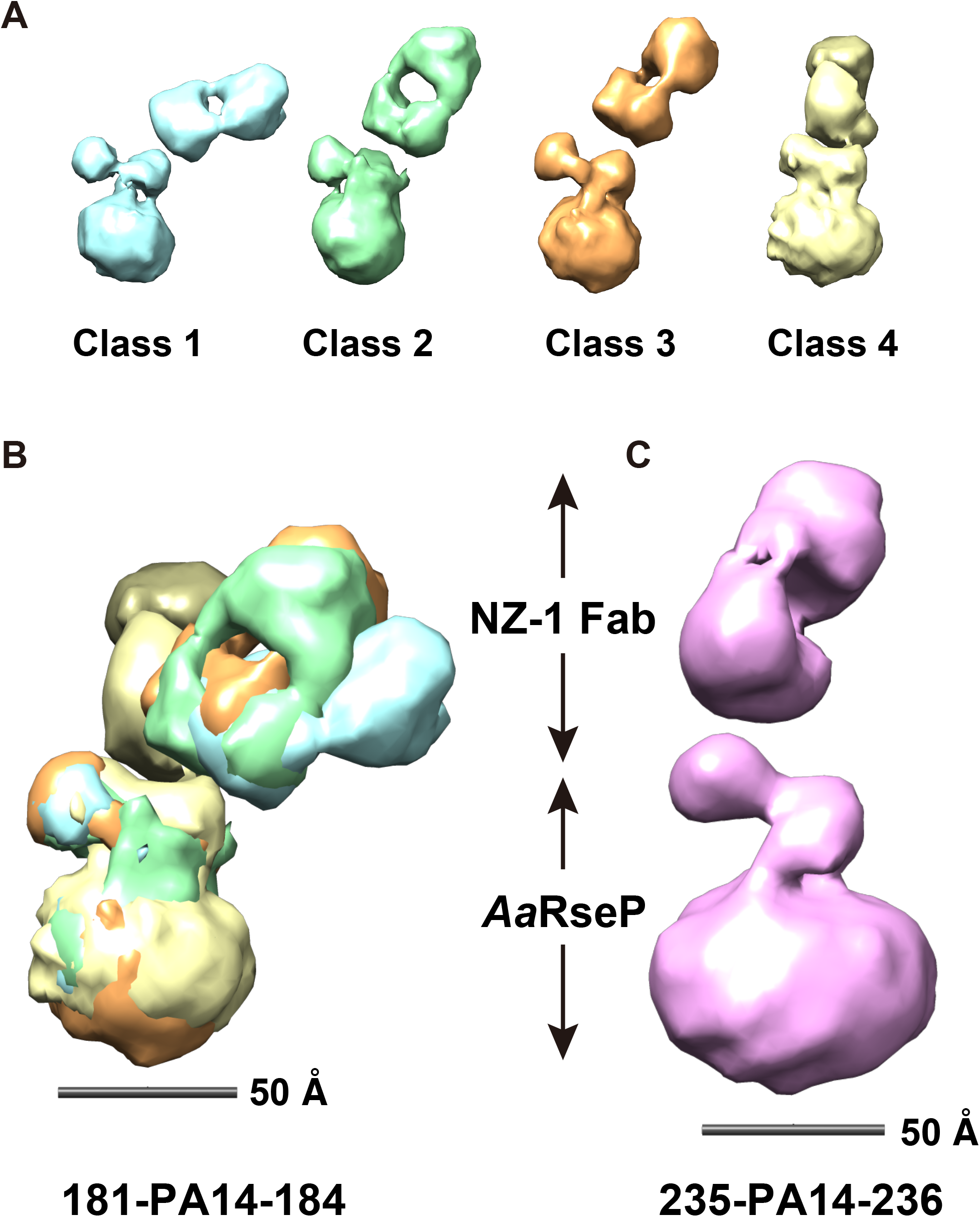
Antibody-assisted negative-staining EM analysis of the PA14-inserted full-length *Aa*RseP. (A) 3D reconstruction models of *Aa*RseP (181-PA14-184) complexed with the NZ-1 Fab. Each of four different 3D reconstructions from the 2D class average images is shown in different colors. (B) Superposition of the 3D reconstructions. The four models in (A) were aligned based on the putative *Aa*RseP region. The NZ-1 Fabs appears to contact *Aa*RseP at almost the same point in each model. (C) 3D reconstruction model of *Aa*RseP (235-PA14-236) complexed with the NZ-1 Fab. The most well-averaged class is drawn as a representative.

### Approximation of the domain arrangement in the full-length *Aa*RseP

We then attempted to superpose the representative models of the Fab-PDZ complexes obtained from the MD simulation onto the 3D reconstruction models of the full-length *Aa*RseP with the NZ-1 Fab from the EM analysis. To select the conformation that appeared with high probability as a representative conformation, we performed the principal component analysis (PCA) for the snapshots within the MD trajectories. For the 181-PA14-184 mutant, the representative model obtained from the PCA (605-ns snapshot in Fig. 5D, Supplementary Fig. 5A) fit the class 1 of the 3D reconstruction model best. Aligning the representative model and the 3D reconstruction at the Fv region resulted in good overlap between the PDZ-N model and one of the protruding lobes of the EM map (Fig. 8A).

**Figure 8.**
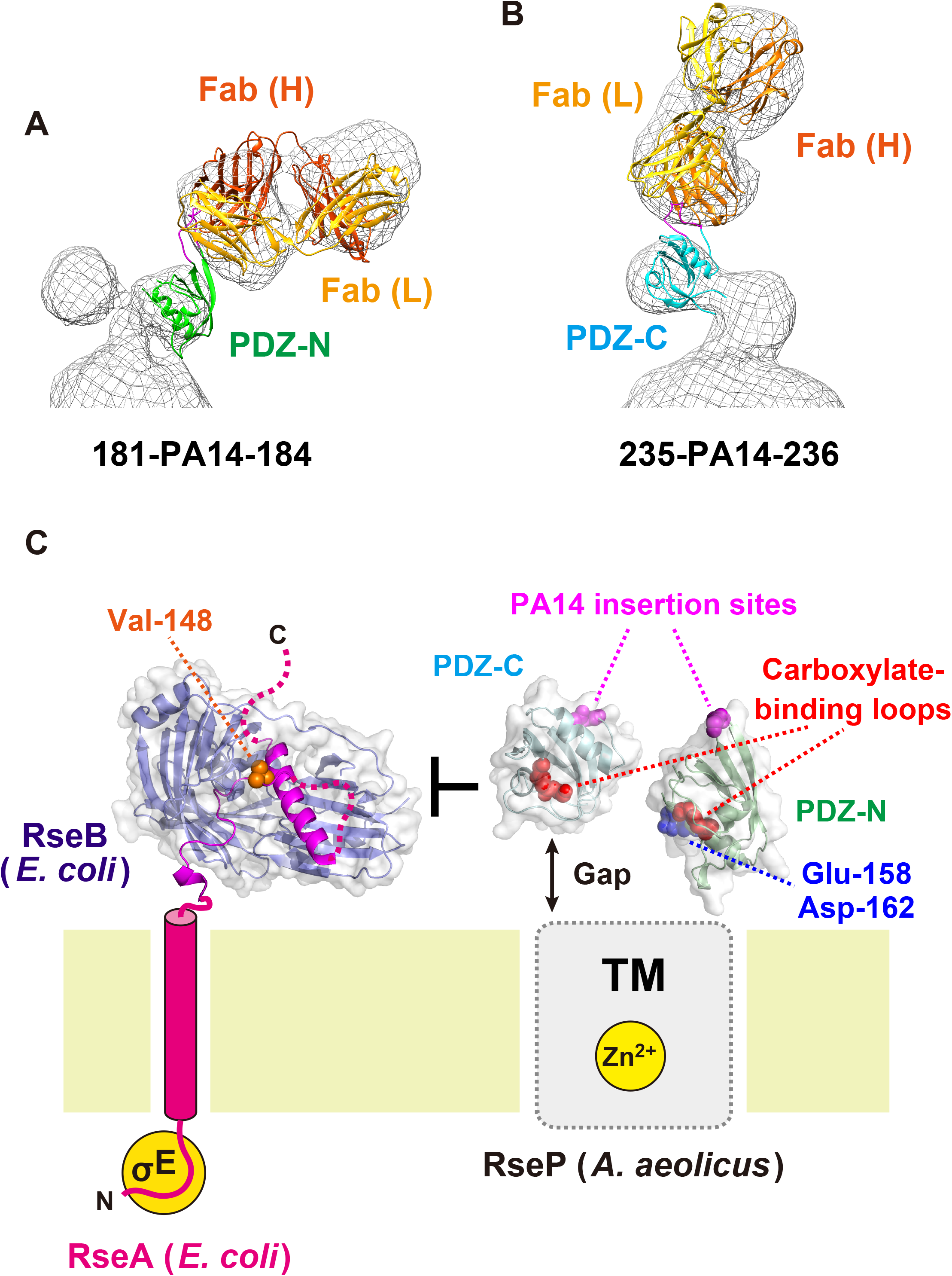
Estimation of the arrangement of two PDZ domains in the full-length *Aa*RseP. (A, B) Structural alignment of the Fab-PDZ models onto the 3D reconstruction. (A) The 605-ns snapshot on the MD trajectory of the NZ-1 Fab-PDZ-N (181-PA14-184) pair, whose conformation was most frequently sampled during the MD simulation, was fit into the 3D reconstruction model of the class 1 shown in Fig. 7A based on the position of the Fv region. (B) The 347-ns snapshot, which showed a similar conformation to that of the complex #2 structure, on the MD trajectory calculated from the complex #1 structure of the NZ-1 Fab-PDZ-C (235-PA14-236) pair was fit into the 3D reconstruction model shown in Fig. 7C based on the position of the Fv region. (C) Structural model of size-exclusion. The approximated arrangement of the two PDZ domains from *Aa*RseP is shown together with the model of RseA from *E. coli* to help understand the size-exclusion mechanism. The crystal structure of RseA periplasmic domain in complex with RseB (PDB ID: 3M4W) is drawn in the RseA model. RseB is shown as blue ribbon model with partial transparent surface. The RseA periplasmic domain is shown as magenta ribbon model, and the side chain of Val-148 is highlighted with orange sphere model. The atomic model of the PDZ tandem from *Aa*RseP was built based on the structural alignments in panels (A, B). First, the two 3D reconstruction models were aligned based on the *Aa*RseP molecule composed of a sphere and two protrusions. Next, the position of the PA14-inserted PDZ-N and -C domains were aligned onto the respective 3D models as described in (A, B). Finally, the crystal structures of wild-type PDZ-N and -C domains were independently aligned onto the models of PA14-inserted mutants. The aligned models of PDZ-N and -C were merged to build the entire PDZ tandem model. The PA14 insertion sites and the carboxylate-binding loops within the ligand-binding grooves are highlighted with magenta and red sphere models, respectively. Glu-158 and Asp-162, shown as blue sphere models, were modeled to be close to the transmembrane region based on the structural alignment. The corresponding residues in *Ec*RseP were estimated to be membrane-proximal residues based on protection from chemical modification. In both 3D reconstruction models, the PDZ-C domain was separated from the TM domain with a gap, which might suppress the entry of the RseB-bound full-length RseA to the active center as the size-exclusion filter.

For the 235-PA14-236 mutant, we first attempted to perform a structural alignment onto the 3D reconstruction using the representative models of the PDZ-C domain complexed with the NZ-1 Fab obtained from the PCA (Supplementary Figs. 3A, 5B). However, neither the representative models calculated from complex #1 nor #2 fit the 3D reconstruction well. Superposition of the NZ-1 Fab region positioned the βA-βB loop of the PDZ-C domain outside the EM map in both two models (Supplementary Fig. 6A, C). These two models adopted conformations similar to that in complex #1 from the crystal structure. In fact, the atomic model of complex #1 also fit the 3D reconstruction poorly (Supplementary Fig. 6A, C). The PDZ-C domain in complex #1 was located closer to the NZ-1 Fab than that in complex #2. In most trajectory snapshots, the PDZ-C domains were also located relatively close to the NZ-1 Fab, as observed in complex #1 (Supplementary Fig. 1B, C). In contrast, the PDZ-C domain in complex #2 was relatively separated from the NZ-1 Fab, and the structural alignment of complex #2 onto the 3D reconstruction resulted in better overlap of the PDZ-C domain with the protruding lobe in the EM map (Supplementary Fig. 6B, D). Within both trajectories initialized from complex #1 and #2, there were minor populations of the snapshots with conformations similar to that in complex #2 (Supplementary Fig. 5B). In those snapshots, the PDZ-C domains were relatively separated from the NZ-1 Fab. In addition, comparison of snapshot models indicated that Lys-255 on PDZ-C and Asp-72 on the NZ-1 light chain were not close enough to form a hydrogen bond in the conformations similar to that in complex #2. In fact, the RMSD of the snapshot model relative to complex #2 showed a negative correlation with the distance between Lys-255 and Asp-72 (Supplementary Fig. 5C). To analyze the structural features of the MD models that fit the 3D reconstruction, we selected one snapshot that had both a low RMSD relative to complex #2 and a large distance between Lys-255 and Asp-72 for each of the trajectories initialized from complex #1 and #2 (Supplementary Figs. 3B, and 5C). According to the PC surface, the selected snapshots were intermediate conformations between complex #1 and #2 (Supplementary Fig. 5B). As a result, both of the two selected MD models (the 347-ns snapshot from complex#1 and the 689-ns snapshot from complex #2) seemed to fit the density similarly to complex #2 (Fig. 8B, and Supplementary Fig. 6B, D). From these observations, we concluded that the hydrogen bond between Lys-255 on PDZ-C and Asp-72 on the NZ-1 light chain is broken in the 3D reconstruction model of the complex.

Finally, combination of the two EM reconstructions enabled us to approximate the spatial arrangement of the PDZ tandem in the context of the full-length protein (Fig. 8C). First, the two EM maps were aligned based on the position of the spherical part and two protrusions. Next, the structural alignments of the MD models onto the EM maps were performed for Fab-PDZ-N complex (the 605-ns snapshot) and Fab-PDZ-C complex (the 347-ns snapshot from complex #1), respectively. Subsequently, the crystal structure of the wild type PDZ-N and C domains were aligned onto the MD models and merged to build the entire PDZ tandem model. In the resultant model, the putative ligand-binding grooves of both two PDZ domains pointed toward the TM domain containing the active center. In addition, both of the 3D reconstruction models commonly indicated that a gap was present between the PDZ-C domain and the TM domain whereas PDZ-N was flush with the spherical profile of the detergent-solubilized TM domain.

## Discussion

Antibody labeling has been widely conducted in structural analysis of protein molecules and complexes. The use of inserted exogenous epitope has the potential to expand its applicability even in case that no antibodies have been established for the target protein. In fact, antibody labeling methods for membrane proteins mediated by the fusion with a thermostabilized variant of apocytochrome *b*_562_ have been reported very recently (Miyagi *et al*., 2020, Mukherjee *et al*., 2020). Our method provides a new option for antibody labeling through exogenous epitopes. However, the structural integrity of the target protein must be maintained for the resulting structural model to be relevant in any of the methods. In our previous study on antibody-assisted structural analysis with NZ-1 labeling, the inserted PA12 tag caused a large structural change around the insertion site in the target protein. In the present study, we attempted to circumvent the deformation that PA12 caused by utilizing the PA14 tag that possesses a pair of Glu-Gly residues upstream of PA12. We anticipated that the two additional residues would act as a buffer region to suppress the structural change in the target. We observed that the two different PA14-inserted PDZ tandem mutants both produced co-crystals with the NZ-1 Fab, which indicated that stable antibody-target complexes were successfully formed through the inserted PA14 tag. In those crystal structures, the PA14 tags were inserted into the two different β-hairpins and adopted closed ring-like conformations when recognized by the NZ-1 Fab in both cases. In these conformations, both ends of the PA14 tag were located less than 7 Å apart based on the Cα-Cα distance of Glu-1’ and Val-14’. Furthermore, there were no significant structural changes observed in the target PDZ domains in comparison to the wild-type structure without the PA14 insertion. These observations suggest that the PA14 tag is suitable both for the insertion into the target and for the complex formation with NZ-1 as compared to the PA12 tag used previously, and open the possibility that the PA14 tag can be inserted into other proteins with varying structural features.

In the present study, we also demonstrated that NZ-1 labeling via an inserted PA14-tag can be applied to negative-stain EM for structural analyses, as demonstrated using the PDZ tandem in *Aa*RseP. As the PDZ tandem is thought to play a pivotal role in the substrate discrimination in RseP, structural data about the spatial arrangement of the PDZ domains should help delineate the regulatory mechanism of intramembrane proteolysis. Our previous chemical modification analysis has suggested that the putative ligand-binding grooves of the two PDZ domains form a pocket-like space sitting just above the RseP active center that is sequestrated within the membrane (Hizukuri *et al*., 2014). The spatial arrangement approximated by the EM analysis is fully consistent with the modification analysis, wherein the putative ligand-binding grooves indeed pointed toward the TM region. In the chemical modification of *Ec*RseP, Asp-162 and Leu-167 on the αB helix were assigned as membrane-proximal residues. In the PDZ tandem model of *Aa*RseP, the corresponding residues, Glu-158 and Asp-162, were located at the interface to the TM domain (Fig. 8C). As the structural changes of the PDZ domains were suppressed by the use of PA14 insertion, the arrangement of the PDZ tandem was thought to be approximated with high fidelity and reflect to the native state. The biochemical analysis confirmed that at least the PA14-inserted mutants showed similar properties to the wild type in the proteolytic reaction within the cell membrane. In our previous study, we proposed that the pocket-forming PDZ tandem serves as a size-exclusion filter that only allows RseA that has been pre-cleaved by DegS to be further cleaved by RseP in *E. coli* (Hizukuri *et al*., 2014). Based on the 3D reconstruction model, the PDZ-C domain was separated from the TM domain. The degree of separation may control substrate discrimination (Fig. 8C). The structural data from our present hybrid analysis will advance biochemical analyses in the investigation of the “size-exclusion model”.

The present study has raised a new issue to be solved when designing PA14-insertion sites for EM structural analysis assisted by NZ-1 labeling. While antibody labeling has application in negative-stain EM (Boisset *et al*., 1993, Boisset *et al*., 1995), the impact of this method has been particularly pronounced on 3D structure determination by cryo-EM. Although the cryo-EM techniques have been drastically improved through the “resolution revolution” of recent years (Kühlbrandt, 2014), it is still difficult to determine high-resolution structures of small proteins due to weak signal from individual particles in the electron micrographs (Rubinstein, 2007). Antibody labeling is expected to resolve this size-limitation problem both by increasing the particle size and by facilitating accurate image alignment (Wu *et al*., 2012), which has already exemplified by recent cryo-EM structure determinations (Kim *et al*., 2019, Coleman *et al*., 2019). Similarly to x-ray crystallography, more rigid complexes with Fabs are, however, desirable to determine high-resolution structures by cryo-EM. While lattice contacts could stabilize the arrangement of proteins in a complex to some extent for x-ray crystallography, individual complex particles in EM micrographs may display more structural heterogeneity. In fact, our negative-stain EM analysis indicated that the orientation of the bound NZ-1 Fab relative to the target PDZ domains was variable. In particular, *Aa*RseP (181-PA14-184) complexed with the NZ-1 Fab showed a high degree of conformational flexibility between the two molecules. Crystallographic analysis showed that only the junction residues of the target PDZ-N domain made direct contacts with the NZ-1 Fab in the 181-PA14-184 mutant. In addition, the MD simulations indicated that the inter-molecular interaction between Glu-1’ on PA14 and NZ-1 can be readily broken in this mutant. For the 181-PA14-184 mutant, we attempted to optimize the insertion site by deleting the turn residues to eliminate the inter-molecular space between the target PDZ-N domain and the bound NZ-1 Fab as that insertion site protruded from the globular domain. Nevertheless, it seems that the conformational flexibility might not be suppressed sufficiently (Fig. 7B) due to the absence of specific interactions between the NZ-1 Fab and the target PDZ-N domain other than through the inserted PA14 tag. In contrast, the orientation of the NZ-1 Fab seemed to be relatively fixed in the complex with *Aa*RseP (235-PA14-236). Lys-255 on PDZ-C interacted with Asp-72 on the NZ-1 light chain in complex #1 of the crystal structure of the PDZ tandem (235-PA14-236) with the NZ-1 Fab, and this interaction was reproduced in the MD simulation. We therefore hypothesized that it could stabilize the subunit arrangements in the complex. However, the structural alignment suggested that only the PA14 residues were involved in the inter-molecular interactions with the NZ-1 Fab because the interaction mediated by Lys-255 was broken in the atomic models that fit the 3D reconstruction from the EM analysis (Supplementary Figs. 3B and 6B, D). As the PA14 tag was inserted into a loop that packs against a flat surface on this insertion mutant, the orientation of the NZ-1 Fab in the EM model might be locked in position by steric hindrance from the flat surface of the PDZ domain. Based on these observations, we conclude that creating an NZ-1-labeled complex with reduced flexibility will require further optimization of the insertion strategy by considering both specific interactions and steric hindrance between the target and the Fab. Computational approaches such as the modelling of PA14-inserted mutant structures and the MD simulation to assess the stabilities of their complexes with the NZ-1 Fab will aid these future optimizations.

In conclusion, we have optimized the insertion method of the NZ-1 epitope to suppress structural changes in the target protein. Using a target of known structure, we showed that inserting the PA14 tag is more suitable for NZ-1 labeling as compared to the PA12 insertion. The advantage of the PA14 tag is that it can reduce structural changes in the target protein during labeling and structure determination even when it was inserted into sterically hindered sites such as loops that pack against the protein. If these advantages are general properties of PA14, the use of PA14 as an inserted epitope tag will expand the range of application of the NZ-1 labeling technique. To further advance our generalizable method of antibody-assisted structural analysis for application in EM studies, future work will optimize the insertion site to fix the orientation of the NZ-1 Fab relative to the target and minimize conformational variability. Other optimization challenges, such as Fab dissociation upon vitrification will also need to be addressed. This endeavor may require mutations of both epitope and paratope. Ultimately, a generalizable method for antibody labeling that produces stable and rigid antibody-target complexes will be an important new tool for high-resolution structural analysis by both x-ray crystallography and cryo-EM.

## Materials and Methods

### Plasmid construction for expression and in vivo cleavage assay

The pGEX-2T-based plasmid for the PDZ tandem fragment (residues 115-292), which was constructed in our previous study (Hizukuri *et al*., 2014), is hereafter termed as pNO1499. The expression plasmids for the PA14-inserted PDZ tandem mutants were constructed by inverse PCR on pNO1499 using primers containing the respective mutations. The PCR products were transformed into *E. coli* XL-1 Blue after digestion of the template pNO1499 with *Dpn*I. The resultant plasmids for the PDZ tandem (181-PA24-184) and (235-PA14-236) mutants are pNY1493, and pNY1468, respectively.

The DNA encoding the full-length *Aa*RseP fused with the C-terminal tag, Gly-Arg-Gly-Ser-His×8 (*Aa*RseP-His_8_) was amplified by PCR using the genomic DNA of *A. aeolicus* VF5 strain and primers encoding the C-terminal tag sequence. The amplified DNA was first cloned into the NdeI/BamHI site of a pET-11c plasmid. Subsequently, the DNA encoding *Aa*RseP-His_8_ together with the Shine-Dalgarno sequence was extracted and cloned into the EcoRI/BamHI site of the plasmid pUC118, resulting in plasmid pNO1457. The expression plasmids for the *Aa*RseP mutants, pNO1461 (active site mutant E18Q), pNY1493 (181-PA14-184), and pNY1478 (235-PA14-236), were constructed by introducing the respective mutations into pNO1457 using the inverse PCR protocol. pTM748 (*Aa*RseP-His_8_), pTM749 (active site mutant E18Q), pTM750 (181-PA14-184), and pTM751 (235-PA14-236) were constructed by introducing a 1.5 kb fragment of EcoRI/BamHI digested pUC118-based plasmids (see above) into the same cloning site on pTWV228. All of these plasmids used in this study were listed in Supplementary Table 1.

### Purification of the PA14-inserted PDZ tandem mutants in complex with the NZ-1 Fab

By using the plasmids constructed above, the PA14-inserted PDZ tandem fragments were produced as N-terminal GST (glutathione S transferase) -fusion proteins, in which a TEV protease recognition site was incorporated between the GST and PDZ tandem sequences, as reported previously (Hizukuri *et al*., 2014). Each PA-inserted construct was overproduced in *E. coli* BL21(DE3) cells and purified from the cell lysate using Glutathione-Sepharose 4B resin (Cytiva, Tokyo, Japan (GE Healthcare)). The mutant fragment was cleaved off from the GST portion through on-column digestion with TEV protease, and the released fragment, which contained two additional residues (Gly-Ser) upstream of the PDZ tandem, was further purified using cation-exchange chromatography HiTrap SP HP (Cytiva, Tokyo, Japan (GE Healthcare)) and size-exclusion chromatography Superdex 200 Increase 10/300 GL (Cytiva, Tokyo, Japan (GE Healthcare)). In parallel, the NZ-1 Fab was prepared by cleaving the NZ-1 antibody using papain and purifying as reported previously (Fujii *et al*., 2016). NZ-1 was obtained from the Antibody Bank (http://www.med-tohoku-antibody.com/topics/antibody.htm) at Tohoku University (Miyagi, Japan). The purified PDZ tandem mutant was mixed with the NZ-1 Fab at a molar ratio of 2:1 and was applied to size-exclusion chromatography to fractionate the complex. The final protein sample was concentrated by ultrafiltration.

### Crystallization and data collection

The initial crystallization conditions were searched using the Index™ (Hampton Research) screening kit containing 96 conditions. 0.2 μL each of protein solution and reagents, respectively, were dispensed into 96-well plates using a Gryphon robotic crystallization system (Art Robbins Instruments) and equilibrated against 60 μL of the reservoir solution by the sitting-drop vapor-diffusion method. Diffraction-quality crystals of the PDZ tandem (181-PA14-184) complexed with the NZ-1 Fab were generated from a crystallization buffer containing 20% (wt./vol.) polyethylene glycol (PEG) 3350, 0.2 M potassium sodium tartrate. Diffraction-quality crystals of the PDZ tandem (235-PA14-236) complexed with the NZ-1 Fab were obtained from a crystallization buffer containing 10%(wt./vol.) PEG 3350, 0.2 M L-proline, and 0.1 M HEPES-Na (pH 7.5). For each crystallization condition, cryoprotectant was prepared by mixing the crystallization buffer and ethylene glycol with a volume ratio of 4:1. All of the crystals were quickly soaked in the cryoprotectant and frozen in liquid nitrogen. X-ray diffraction data were collected using a photon counting pixel array detector PILATUS3 S 6M (Dectris) at Photon Factory (PF) BL-5A and 17A (Tsukuba, Japan). Data were processed and scaled with XDS (Kabsch, 2010) and aimless (Evans & Murshudov, 2013). Diffraction intensities were converted to structure factors using the CCP4 programs where 5% of the unique reflections were randomly selected as a test set for the calculation of free *R*-factor (Winn *et al*., 2011). Data collection statistics are summarized in Table 1.

### Crystallographic analysis

For both of the co-crystals, initial phases were determined by the molecular replacement method by using Molrep (Vagin & Teplyakov, 1997) in CCP4. First, the Fv and constant regions of the NZ-1 Fab were assigned separately using the atomic coordinates of the NZ-1 Fab bound to the PA14 peptide (Fujii *et al*., 2016) (PDB code: 4YO0). Next, the PDZ-N and -C domains were searched separately using the atomic coordinates of the *A. aeolicus* PDZ tandem (Hizukuri *et al*., 2014) (PDB code: 3WKL) with the NZ-1 Fab models fixed in the asymmetric unit. The assigned models were manually fit into the electron density map using the program COOT (Emsley *et al*., 2010). The updated models were refined with phenix.refine (Afonine *et al*., 2012) iteratively while monitoring the stereochemistry with MolProbity (Chen *et al*., 2010). As reported in the previous study (Tamura *et al*., 2019), the electron densities indicated that Asn-150 side chain reacted with the main chain amide group of Gly-151 and formed a succinimide (Snn) group in the PDZ tandem (181-PA14-184) mutant. Refinement statistics are summarized in Table 2. The atomic coordinates of the Fab-complexes of the PDZ tandems (181-PA14-184) and (235-PA14-236) were deposited in the Protein Data Bank with the accession codes, 7CQC and 7CQD, respectively. Structural superposition and RMSD calculation were performed by the pair-wise alignment protocol using LSQKAB (Kabsch, 1976). Figures for protein structures were prepared with PyMOL (The PyMOL Molecular Graphics System, Version 2.3 Schrödinger, LLC.).

### In vivo *cleavage assay of* Aa*RseP and the mutants*

The *in vivo* proteolytic activity of *Aa*RseP was analyzed using *E. coli* KK211 (Δ*rseA*, Δ*rseP*) cells (Kanehara *et al*., 2002) as described previously (Akiyama *et al*., 2015, Hizukuri *et al*., 2017). *E. coli* KK211 (Δ*rseA*, Δ*rseP*) cells harboring pYH124 (HA-MBP-RseA(LY1)148) were transformed with a plasmid pKK11 (*Ec*RseP-His_6_-Myc) (Kanehara *et al*., 2001), pTM748 (*Aa*RseP-His_8_), pTWV228 or plasmids encoding their derivatives. M9 medium (without CaCl_2_) (Miller, 1972) supplemented with 20 µg/mL of each of the 20 amino acids, 2 µg/mL thiamine, 0.4% glucose, 1 mM IPTG and 5 mM cAMP was inoculated with transformed *E. coli* KK211 cells and grown at 30°C for 3 hours. Proteins were precipitated by trichloroacetic acid (TCA) treatment and separated by Laemmli SDS-PAGE. Immunoblots with anti-HA, anti-His or anti-SecB antibodies were visualized by Lumino image analyzer LAS-4000 mini (Cytiva, Tokyo, Japan (GE Healthcare)) using ECL Prime Western Blotting Detection Reagents (Cytiva, Tokyo, Japan (GE Healthcare)). Rabbit polyclonal anti-HA (HA-probe (Y-11), Santa Cruz Biotechnology) and anti-SecB (a gift from Shoji Mizushima’s Lab.) antibodies were used for immunoblotting. For detection of His-tagged proteins, anti-His antibodies from the Penta-His HRP Conjugate Kit (Qiagen) were used.

### *Purification of the* Aa*RseP mutants and complex formation with the NZ-1 Fab*

For the EM analysis, the PA14-inserted *Aa*RseP mutant was overproduced in *E. coli* KK374 (Δ*rseA*, Δ*rseP*, Δ*degS*) (Akiyama *et al*., 2004). *E. coli* KK374 cells transformed with the expression plasmids (Table 1) were grown at 30°C to an OD_600_ of 0.7 in a medium containing 10 g of bactotryptone, 5 g of yeast extract and 10 g of NaCl per liter supplemented with 50 μg/mL ampicillin, followed by induction of overexpression with 0.1 mM IPTG and incubation at 30°C for additional 4 hours. Cells were harvested by centrifugation and lysed by sonication in 10 mM Tris-HCl (pH 7.4), 150 mM NaCl. The cell lysates were centrifuged at 40,000 ×g for 45 min. at 277 K. Subsequently, the supernatant was further separated by ultracentrifuge at 200,000 ×g for 90 min at 277 K. The membrane fraction collected as a precipitant was suspended in 10 mM Tris-HCl (pH 7.4), 150 mM NaCl and was ultra-centrifuged again under the same conditions. Finally, the precipitated membrane fraction was suspended in 10 mM Tris-Cl, 150 mM NaCl and the total protein was quantified using the bicinchoninic acid (BCA) assay. The suspension of the membrane fraction was diluted with the same buffer to adjust the protein concentration to 10 mg/mL using bovine serum albumin as a standard.

Membrane proteins were solubilized by adding the same volume of a solubilization buffer containing 40 mM Tri-HCl (pH 8.0), 150 mM NaCl, and 2% n-dodecyl-N,N-dimethylamine-N-oxide (DDAO) to the above-prepared suspension of the membrane fraction. After incubation at 277 K for 1 hour, the mixture was ultra-centrifuged at 210,000 ×g for 90 min. at 277 K. The supernatant was applied to Ni-NTA agarose resin and the unbound fraction was washed out with a buffer containing 20 mM Tris-HCl (pH 8.0), 300 mM NaCl, 20 mM imidazole, and 0.05% DDAO. The resin was further washed with a buffer containing 20 mM Tris-HCl (pH 8.0), 300 mM NaCl, 50 mM imidazole, and 0.1% GDN for detergent exchange. The *Aa*RseP mutant was eluted from the resin with a buffer containing 20 mM Tris-HCl (pH 8.0), 300 mM NaCl, 250 mM imidazole, and 0.1% GDN. The eluted mutant was then applied to a Superdex 200 10/300 GL size-exclusion chromatography column (Cytiva, Tokyo, Japan (GE Healthcare)) to isolate the monodisperse fraction of the *Aa*RseP mutant. Finally, the purified *Aa*RseP mutant was mixed with the NZ-1 Fab and was again applied to the size-exclusion chromatography to separate the complex fraction.

### Negative-stain electron microscopy

All purified samples were diluted to 1.0 μg/mL. For negative-stain EM, 5 μL of protein was applied to glow-discharged, 600 mesh, carbon-coated grids. The grids were negatively stained with 2% ammonium molybdate, blotted with filter paper, and air dried. JEM2200FS (JEOL) was operated at 200 kV to acquire micrographs using a K2 camera (Gatan) in counting mode. Micrographs were recorded at a magnification of 20,000x (1.98Å/pix) with defocus of −0.5 to −2.0 μm. A movie of 50 frames was taken for each micrograph, and motion correction was performed using MotionCor2 (Zheng *et al*., 2017). Contrast transfer function was estimated by Gctf (Zhang, 2016). All subsequent processing was carried out using RELION3 (Zivanov *et al*., 2018). Particles were selected using the Relion Autopicker and three rounds of 2D classification were performed to select particles for 3D reconstruction. Fitting of the atomic coordinates of Fab-PDZ complex into 3D reconstruction model and figure preparation were performed with Chimera (Pettersen *et al*., 2004).

### Molecular dynamics simulation

Initial models were prepared by performing the energy minimization using the crystal structures of the PA14-mediated complexes between the NZ-1 Fab and the PDZ tandems from this study. For both of the two PDZ tandems, only the PDZ domain that contains the PA14 insertion was included in the initial model. For PDZ-N (181-PA14-184), residues 113-206, in which Gly-113 and Ser-114 were derived from the expression tag, were used in the simulation while residues 207-292 were included for the PDZ-C (235-PA14-236) model. The disordered loop regions in the NZ-1 Fab were built by MODELLER (Sali & Blundell, 1993). The succinimide in PDZ-N and the pyroglutamate at the N-terminus of the NZ-1 light chain were remodeled as aspartate and glycine, and as glutamate, respectively. The solvent system around each Fab-PDZ complex was prepared by the Solution Builder as implemented in CHARMM-GUI (Jo *et al*., 2008, Lee *et al*., 2016). The protonation state of histidine was calculated by PROPKA (Olsson *et al*., 2011, Søndergaard *et al*., 2011) implemented in PDB2PQR (Dolinsky *et al*., 2004) at pH 7. Missing hydrogen atoms were inserted using the Solution Builder, and the N- and C-termini were set to NH_3+_ and COO^-^, respectively. The MD unit cell was set to a rectangular cell with a minimum of 10 Å edge distance between the protein and the walls of the cell. The cell was filled with the TIP3P water model (Jorgensen *et al*., 1983) with 150 mM NaCl and added Na^+^ counterions.

The all-atom MD simulations were carried out using the GROMACS ver. 2016.3 MD program package (Abraham *et al*., 2015, Pronk *et al*., 2013) with CHARMM36m force field (Huang *et al*., 2017, MacKerell *et al*., 1998, MacKerell *et al*., 2004) under periodic boundary conditions. The electrostatic interactions were handled by the smooth particle mesh Ewald method (Essmann *et al*., 1995), and the van der Waals interactions were truncated by a switching function with a range of 10-12 Å. The bond lengths involving hydrogen atoms were constrained by the P-LINKS algorithm (Hess, 2008). According to the default setup of the CHARMM-GUI, an energy minimization and a 125 ps equilibration run as *NVT* ensemble with a 1 fs timestep were executed before the production run. The production run was performed as *NPT* ensemble with a 2 fs timestep. The Nosé-Hoover scheme was used for the thermostat (Hoover, 1985, Nosé, 1984), and the Parrinello-Rahman approach was used as the barostat (Nosé & Klein, 1983, Parrinello & Rahman, 1981). The temperature and pressure were set to 300 K and 1 atm, respectively. The simulation length of the production run was 1 μs.

RMSD and RMSF values were calculated using snapshots extracted from each 1-μs simulation at every 1 ns (1000 snapshots for each trajectory). For PDZ-N, the calculations were performed over the Cα atoms of the residues 123-206 because the residues 113-122 in the N-terminal region is not included in the PDZ-fold and exhibited large fluctuation during the simulation compared to the remaining region of the PDZ-N domain. To estimate the structural change in the PDZ domains, RMSDs were calculated relative to the energy-minimized initial models of the respective PDZ domains, which were almost identical to the atomic models in the corresponding crystal structures, as shown in Fig. 5A. To estimate the degree of fluctuation in the orientation of the PDZ domains relative to the NZ-1 Fab, RMSFs were calculated over the averaged structures of the respective complexes after aligning the snapshot models based on the V_H_ region (residues 20-130) in Fig. 5B, C. In addition, RMSDs were calculated for the PDZ domains by aligning the snapshot models within a trajectory of a Fab-PDZ complex to the corresponding initial model based on the V_H_ region in Fig. 5E, F and Supplementary Fig. 2. Furthermore, the principal component analysis (PCA) was carried out by using the snapshot models aligned at the V_H_ region to separate out the characteristic movements in the fluctuations between the NZ-1 Fab and the respective PDZ domains. For the NZ-1 Fab-PDZ-C (235-PA14-236) complex, the two trajectories for complex #1 and #2 were merged in the PCA calculation. The structural distribution was represented as a 2D normalized histogram of -ln(Z) plotted on a PC map with a pixel size of 10 Å × 10 Å, where Z was the probability of a given conformation, as shown in Supplementary Fig. 5A, B. Hierarchical clustering was implemented using the MMTSB tool (Feig *et al*., 2004) to select the representative model with the maximum probability for each trajectory: the 605-ns snapshot for the NZ-1 Fab-PDZ-N (181-PA14-184), the 317-ns snapshot for complex #1 of the NZ-1 Fab-PDZ-C (235-PA14-236), and the 444-ns snapshot for complex #2, respectively (Supplementary Fig. 5A, B).

## Acknowledgements

We are grateful to the beamline staff of Photon Factory (Tsukuba, Japan) for providing data collection facilities and support. We thank Samuel Thomson for editing the manuscript, and prof. Junichi Takagi for useful discussion. This research is partially supported by the Japan Society for the Promotion of Science (JSPS) KAKENHI under Grant Numbers JP26291016, JP17K19206, and 19H03170 (to T.N.), under JP19K06562 (to Y.H.), and under JP18H023404 (to Y.A.), by Grant-in-Aid for Scientific Research on Innovative Areas from Ministry of Education, Culture, Sports, Science and Technology (MEXT) under Grant Number 18H05426 (to M.I.), by Platform Project for Supporting in Drug Discovery and Life Science Research (Platform for Drug Discovery, Informatics, and Structural Life Science) from the Japan Agency for Medical Research and Development (AMED) under Grant Numbers JP16am0101020 (to T.N.), by the Platform Project for Supporting Drug Discovery and Life Science Research (Basis for Supporting Innovative Drug Discovery and Life Science Research (BINDS)) from AMED under Grant Numbers 19am0101072j0003 (support number 0418) (to M.H., and K.I.), JP20am0101009 (support number 2633) (to M.I.), JP20am0101078 (support number 1948) (to Y.K.), by Program for Promoting Researches on the Supercomputer Fugaku (MD-driven Precision Medicine) under Project ID: hp200129, by RIKEN Dynamic Structural Biology Project (to M.I.), and by AMED under JP20am0401013, and JP20ae0101028 (to Y.K.).

This work was performed in part under the Cooperative Research Program (Joint Usage/Research Center program)of the Institute for Frontier Life and Medical Sciences, Kyoto University, under the Collaborative Research Program of the Institute for Protein Research, Osaka University, CR-19-05, and under the Cooperative Research Project Program of Life Science Center for Survival Dynamics, Tsukuba Advanced Research Alliance (TARA Center), University of Tsukuba.

## Author contributions

T.N. conceived the project. R.T.-S., and T.N. designed the research. R.T.-S., and R.A. prepared the target proteins and performed the crystallographic analysis. M.K.K., and Y.K. produced the antibody. R.O. prepared the antibody fragment for structural analysis. K.I. supervised the EM work, and M.H. obtained and analyzed the EM images. T.M., Y.H., and Y.A. examined the enzymatic activity of the mutants. T.E., and M.I. performed the MD simulations. R.T.-S., R.A., and T.N. analyzed the data from structural and functional analysis. All the authors contributed to paper preparation, and T.N. compiled the paper.

