## Supplementary Information for "Moving toward generalizable NZ-1 labeling for 3D structure determination with optimized epitope tag insertion"

### Supplementary Table 1. Plasmid list

| Plasmid | Vector | Encoded protein | Reference or source |
|---|---|---|---|
| pGEX-2T | | | Cytiva (GE healthcare) |
| pNO1499 | pGEX-2T | PDZ tandem | (Hizukuri *et al.*, 2014) |
| pNY1468 | pGEX-2T | PDZ tandem (235-PA14-236) | This study |
| pNY1493 | pGEX-2T | PDZ tandem (181-PA14-184) | This study |
| pET-11c | | | Agilent Technologies (Stratagene) |
| pUC118 | | | Takara Bio |
| pTWV228 | | | Takara Bio |
| pYH124 | pTWV228 | HA-MBP-RseA(LY1)148 | (Hizukuri & Akiyama, 2012) |
| pKK11 | pTWV228 | *Ec*RseP-$His_6$-Myc | (Kanehara *et al.*, 2001) |
| pKK34 | pTWV228 | *Ec*RseP(E23Q)-$His_6$-Myc | (Kanehara *et al.*, 2001) |
| pNO1457 | pUC118 | *Aa*RseP-$His_8$ | This study |
| pNO1461 | pUC118 | *Aa*RseP(E18Q)-$His_8$ | This study |
| pNY1478 | pUC118 | *Aa*RseP(235-PA14-236)-$His_8$ | This study |
| pNY1498 | pUC118 | *Aa*RseP(181-PA14-184)-$His_8$ | This study |
| pTM748 | pTWV228 | *Aa*RseP-$His_8$ | This study |
| pTM749 | pTWV228 | *Aa*RseP(E18Q)-$His_8$ | This study |
| pTM750 | pTWV228 | *Aa*RseP(235-PA14-236)-$His_8$ | This study |
| pTM751 | pTWV228 | *Aa*RseP(181-PA14-184)-$His_8$ | This study |

### Supplementary Figures

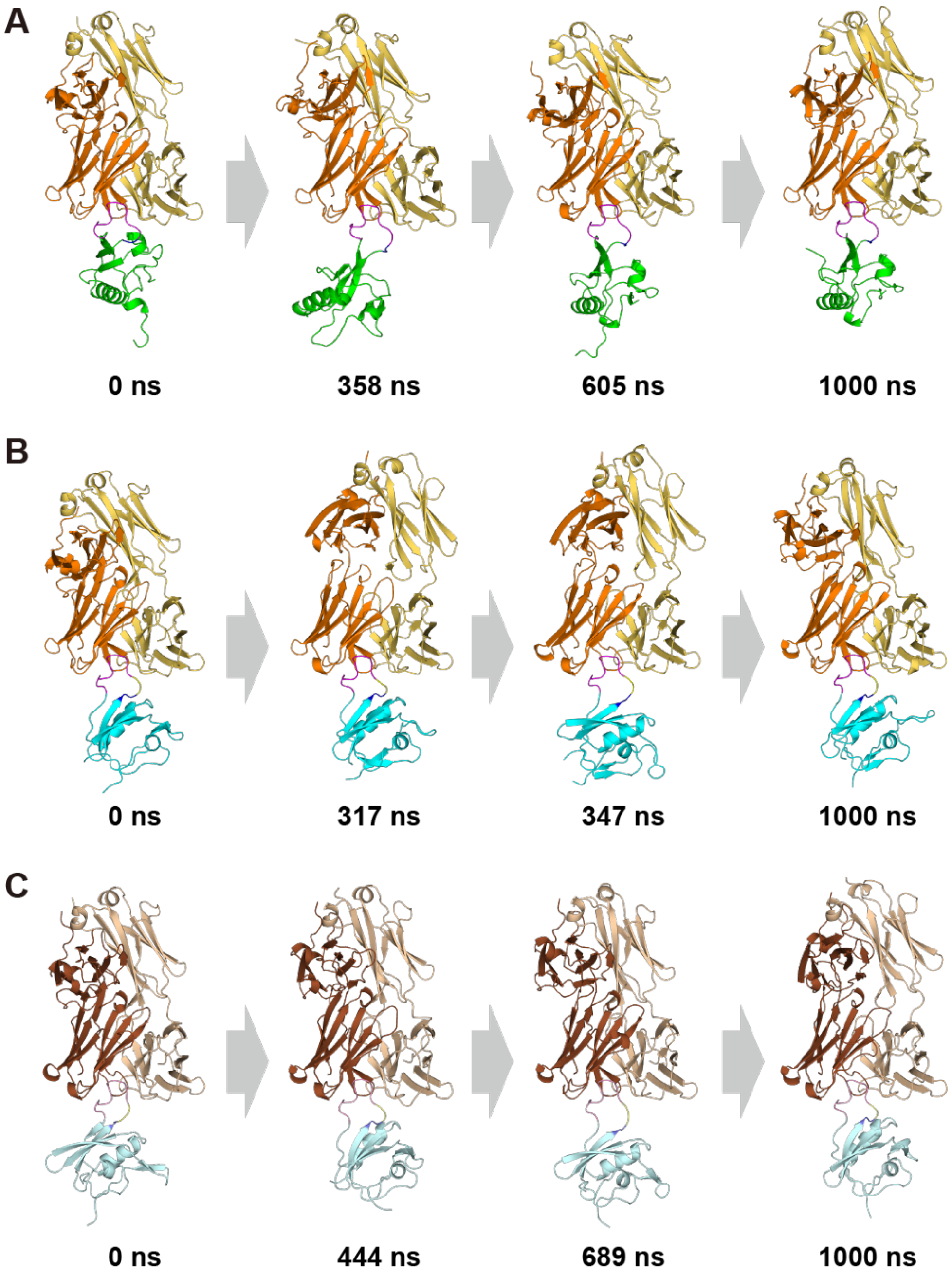


**Supplementary Figure 1**

Snapshot models from the MD trajectories. (A) The trajectory of PDZ-N (181-PA14-184) complexed with the NZ-1 Fab. Snapshots taken during 1-μs simulation time are shown as ribbon models. The heavy and light chain of the NZ-1 Fab are colored dark and light orange, respectively. The PDZ-N domain (residues 113-206) is colored green. The inserted PA14 tag is colored magenta except for Glu-1’ and Gly-2’ colored blue. The snapshot at 0 ns was the energy-minimized initial model of PDZ-N (181-PA14-184) complexed with the NZ-1 Fab while that at 1000 ns was the final snapshot model in the 1-μs simulation. The snapshot at 358 ns showed the highest RMSD relative to the initial model. The snapshot at 605 ns was the representative model selected from the PCA in Supplementary Fig. 5A. (B) The trajectory of PDZ-C (235-PA14-236) complexed with the NZ-1 Fab initialized from the energy-minimized model of complex #1. The NZ-1 Fab is colored as in the panel (A). The PDZ-C domain (207-292) is colored cyan. For the inserted PA14 tag, Glu-1’ and Gly-2’ are colored blue while Gly-3’ and Val-4’ are colored yellow. The remaining 10 residues of the PA14 tag are colored magenta. The snapshot at 317 ns was the representative model selected from the PCA in Supplementary Fig. 5B. The snapshot at 347 ns was the model showing both the low RMSD relative to the PDZ-C model in complex #2 and the long distance between Lys-255 on PDZ-C and Asp-72 on the NZ-1 light chain in Supplementary Fig. 5C. (C) The trajectory of PDZ-C (235-PA14-236) complexed with the NZ-1 Fab initialized from the energy-minimized model of complex #2. The heavy and light chain of the NZ-1 Fab are colored dark and light brown, respectively. For the inserted PA14 tag, Glu-1’ and Gly-2’ are colored light blue while Gly-3’ and Val-4’ are colored light yellow. The remaining 10 residues of PA14 are colored light magenta. The snapshot at 444 ns was the representative model selected from the PCA in Supplementary Fig. 5B. The snapshot at 689 ns was the model showing both the low RMSD relative to the PDZ-C model in complex #2 and the long distance between Lys-255 on PDZ-C and Asp-72 on the NZ-1 light chain in Supplementary Fig. 5C.

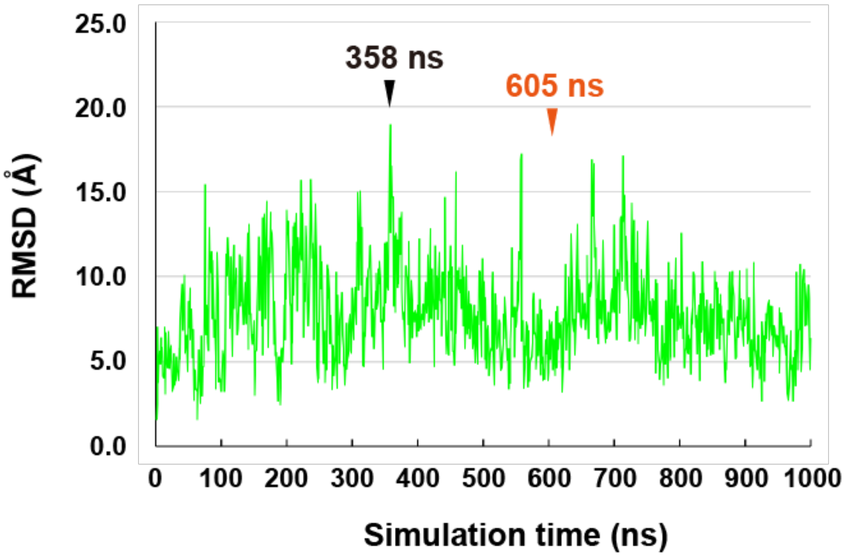


**Supplementary Figure 2**

Deviation of the position of the PDZ-N domains relative to the NZ-1 Fab in the MD simulations. The snapshot models were aligned to the initial model, which was constructed by the energy minimization of the crystal structure of PDZ-N (181-PA14-184) complexed with the NZ-1 Fab, based on the $V_H$ region. Subsequently, the RMSDs for the PDZ-N domains were calculated between the initial model and the respective snapshots on the trajectory. PDZ-N (181-PA14-184) showed the highest RMSD value relative to the initial model at 358 ns. The snapshot at 605 ns was selected as the representative model from the PCA in Supplementary Fig. 5A.

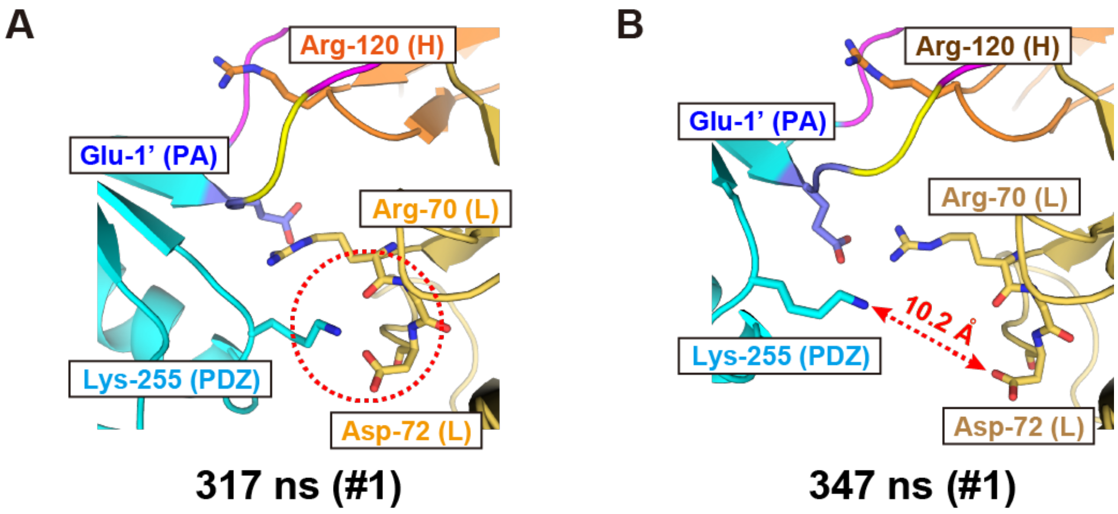


**Supplementary Figure 3**

Binding interfaces of the snapshot models of PDZ-C (235-PA14-236) complexed with the NZ-1 Fab. (A) Residues at the binding interface in the 317-ns snapshot within the trajectory initialized from complex #1. Lys-255 on the PDZ-C domain formed hydrogen bonds with the side chain of Asp-72 and the main chain of Arg-70 on the NZ-1 light chain. These interactions seemed to draw the PDZ-C domain close to the NZ-1 Fab. (B) Residues at the binding interface in the 347-ns snapshot within the trajectory initialized from complex #1. The side chain amino group of Lys-255 on the PDZ-C domain was separated by 10.2 Å from the side chain carboxyl group of Asp-72 on the NZ-1 light chain.

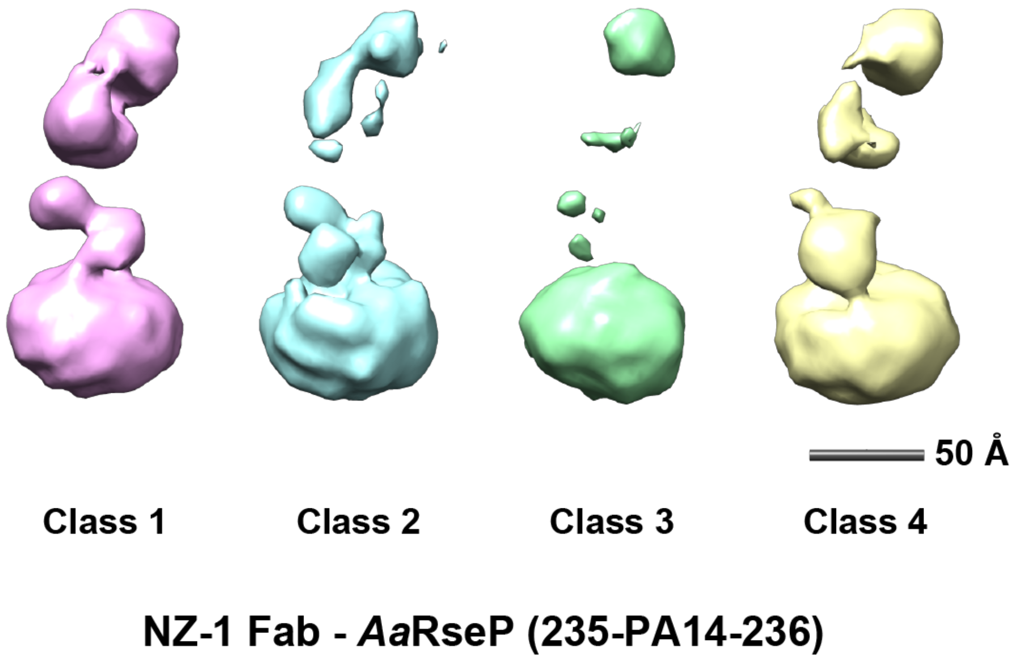


**Supplementary Figure 4**

3D reconstruction models of *Aa*RseP (235-PA14-236) complexed with the NZ-1 Fab. Four different 3D models from the 2D class average images are shown in different colors. Class 1 corresponds to the 3D model shown in Fig. 7C. The 3D models of classes 2 to 4 were aligned onto that of class 1 based on the position of the putative NZ-1 Fab part. The densities for the NZ-1 Fab and the PDZ tandem region were relatively weak and unresolved in classes 2 to 4, as compared to that in class 1.

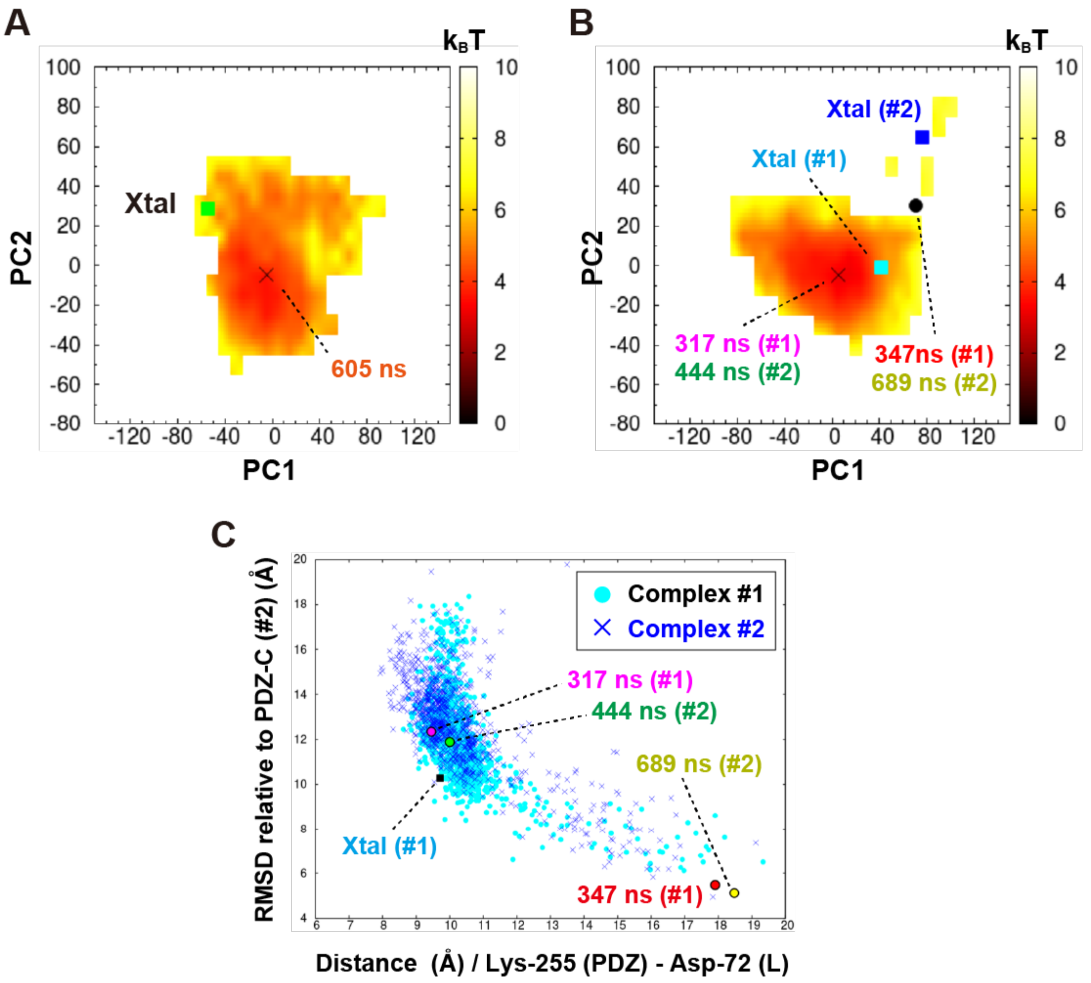


**Supplementary Figure 5**

Analysis of structural fluctuation and domain orientation within the Fab-PDZ complex. (A) Quantification of structural fluctuation of PDZ-N (181-PA14-184) complexed with NZ-1 Fab. PCA was performed for the snapshots on the trajectory to quantify the structural fluctuation. The structural distribution is represented as a 2D normalized histogram of -ln(Z) plotted on a PC map with a pixel size of 10 Å × 10 Å, where Z was the probability of a given conformation. The values for the free energy are given in units of $k_BT$ (color bar), where $k_B$ is the Boltzmann constant and T is the temperature. The initial model of PDZ-N complexed with the NZ-1 Fab (Xtal), which was almost identical to the crystal structure, is indicated with the green square on the plot. The 605-ns snapshot was selected from the cluster with the maximum probability on the PC map. (B) Quantification of structural fluctuation of PDZ-C (235-PA14-236) complexed with the NZ-1 Fab. Snapshots from two trajectories initialized from each of complexes #1 and #2 were merged and subjected to the PCA. The structural distribution is represented as a 2D normalized histogram, as in panel (A). The initial models of NZ-1 Fab-PDZ-C pair constructed from complex #1 and #2, which are almost identical to the corresponding crystal structures, are indicated with a cyan (Xtal (#1)) or blue (Xtal (#2)) squares on the plot. The 317-ns snapshot from complex #1 (magenta) and the 444-ns snapshot from complex #2 (green) were selected from the cluster with the maximum probability on the PC map. The 347-ns snapshot from complex #1 (red) and 689-ns snapshot from complex #2 (yellow), which showed both the low RMSD relative to the PDZ-C model in complex #2 and the long distance between Lys-255 on PDZ-C and Asp-72 on the NZ-1 light chain, are also plotted on the PC map. (C) Analysis of domain orientation within the Fab-PDZ complex based on comparison to complex #2. The RMSD relative to the PDZ-C model in complex #2 for each snapshot was plotted against the distance between the Cα atoms of Lys-255 on PDZ-C and Asp-72 on the NZ-1 light chain. The snapshots from the trajectories for complexes #1 and #2 are indicated with cyan circles and blue cross marks, respectively. In complex #2, the PDZ-C domain was separated from the NZ-1 Fab while the hydrogen bond between Lys-255 and Asp-72 was broken. Therefore, the low RMSD and the long distance between Lys-255 and Asp-72 indicate the high conformational similarity to complex #2. The 317-ns snapshot from complex #1 (magenta) and the 444-ns snapshot from complex #2 (green) were located in proximity of the majority of snapshots on the PC map, whereas the 347-ns snapshot from complex #1 (red) and 689-ns snapshot from complex #2 (yellow) appeared in a minor population of outliers.

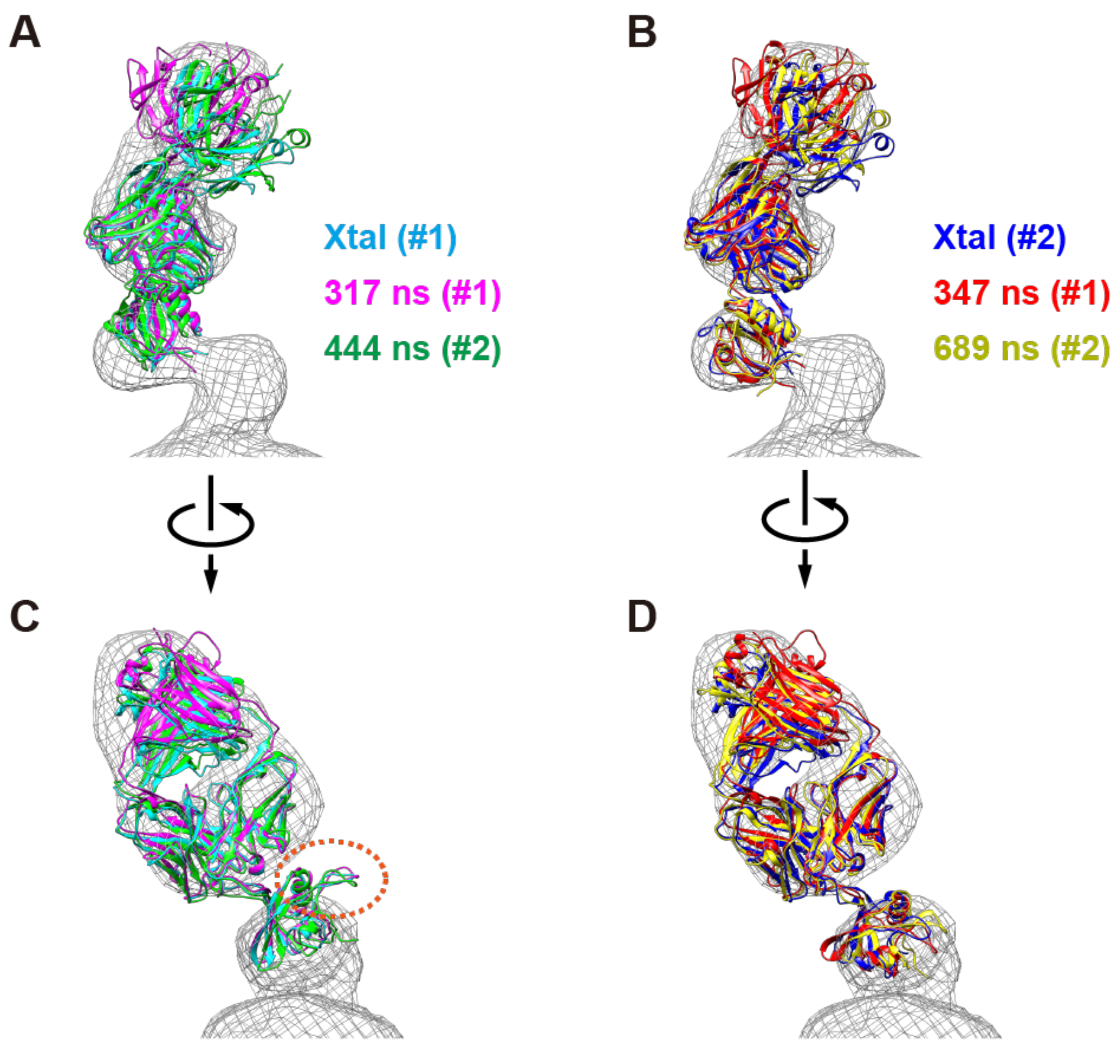


**Supplementary Figure 6**

Structural alignment of the NZ-1 Fab-PDZ-C complexes onto the 3D reconstruction model. (A) Complex #1 (cyan) from the crystal structure of the PDZ tandem (235-PA14-236) complexed with the NZ-1 Fab, the 317-ns snapshot from the MD trajectory initialized from complex #1 (magenta), and the 444-ns snapshot from the trajectory initialized from complex #2 (green) were aligned onto the 3D reconstruction model of *Aa*RseP (235-PA14-236) complexed with the NZ-1 Fab. (B) Complex #2 (blue), the 347-ns snapshot from the trajectory initialized from complex #1 (red), and the 689-ns snapshot from the trajectory initialized from complex #2 (yellow) were aligned onto the 3D reconstruction model. Panels (C) and (D) are side-views of panels (A) and (B), respectively. For the three complexes shown in (A) and (C), alignment of the models based on the position of the Fv region placed the βA-βB loops of the PDZ-C models outside the EM map, as highlighted with red dotted circle, while the same loops on the three models in (B) and (D) largely fit within a lobe of density in the EM map.
